# The independent influences of age and education on functional brain networks and cognition in healthy older adults

**DOI:** 10.1101/154898

**Authors:** Alistair Perry, Wei Wen, Nicole A. Kochan, Anbupalam Thalamuthu, Perminder S. Sachdev, Michael Breakspear

## Abstract

Healthy ageing is accompanied by a constellation of changes in cognitive processes and alterations in functional brain networks. The relationships between brain networks and cognition during ageing in later life are moderated by demographic and environmental factors, such as prior education, in a poorly understood manner. Using multivariate analyses, we identify three latent patterns (or modes) linking resting-state functional connectivity to demographic and cognitive measures in 101 cognitively-normal elders. The first mode (*p*=0.00043) captures an opposing association between age and core cognitive processes such as attention and processing speed on functional connectivity patterns. The functional subnetwork expressed by this mode links bilateral sensorimotor and visual regions through key areas such as the parietal operculum. A strong, independent association between years of education and functional connectivity loads onto a second mode (*p*=0.012), characterised by the involvement of key hub-regions. A third mode (*p*=0.041) captures weak, residual brain-behaviour relations. Our findings suggest that circuits supporting lower-level cognitive processes are most sensitive to the influence of age in healthy older adults. Education, and to a lesser extent, executive functions, load independently onto functional networks - suggesting that the moderating effect of education acts upon networks distinct from those vulnerable with ageing. This has important implications in understanding the contribution of education to cognitive reserve during healthy ageing.

## Introduction

Healthy ageing in the later decades of life is associated with progressive changes in cognition which impact upon on function and inter-personal relationships (Stuck, et al., 1999; Willis, et al., 2006). Fluid-based cognitive functions such as processing speed, executive function and working memory are particularly sensitive to changes that arise from age-related neurobiological processes (Grady, 2012; Park and Reuter-Lorenz, 2009). More rapid morphological changes (indexed by volumetric size and thickness) in pre-frontal, hippocampal, and parietal cortices is thought to underpin progressive age-related cognitive changes (Dennis and Cabeza, 2008; Park and Reuter-Lorenz, 2009; Raz, et al., 2005). However, investigations that have reported morphometric changes associated with age-related variability in cognitive performance are somewhat inconsistent or even contradictory (Rodrigue and Kennedy, 2011). It is crucial to disambiguate the trajectory of normal age-related changes in later life from the pathology of neurodegenerative disorders such as Alzheimer’s disease (AD) (Dennis and Thompson, 2014).

Univariate alterations in morphological brain structures only partially capture the complexity of neurobiological changes associated with ageing (Rodrigue and Kennedy, 2011). Recent network conceptualizations propose that human brain function is shaped by interactions (connections) between its constituent elements (brain regions) through neural networks that possess a complex topological organization (Bassett and Bullmore, 2006; Bullmore and Sporns, 2012; Sporns, 2013). Brain networks delicately balance the opposing requirements for functional integration and segregation, giving rise to complex cognitive and perceptual functions (Friston, et al., 1995; Sporns, et al., 2000; Tononi, et al., 1994). Networks of whole-brain functional connectivity patterns can be constructed from the temporal correlations of spontaneous fluctuations in neurophysiological signals between brain regions, and analysed with graph-theoretical approaches (Biswal, et al., 1995; Fornito, et al., 2013; Fox and Raichle, 2007).

Fluid cognitive functions require patterns of integrated and coordinated neural interactions, suggesting age-related variability in performance may be attributable to corresponding changes in large-scale connectivity (Andrews-Hanna, et al., 2007). Investigations into intrinsic resting-state networks (RSN) (Damoiseaux, et al., 2006; Fox, et al., 2005) have consistently revealed reduced functional connectivity in core cognitive systems such as the Default-Mode Network (DMN) over the lifespan (Damoiseaux, 2017; Damoiseaux, et al., 2008). On the other hand, connectivity between-networks has been found to increase - indicative of decreased functional specialisation with age (Betzel, et al., 2014; Chan, et al., 2014; Ferreira, et al., 2015; Geerligs, et al., 2015; Grady, et al., 2016; Ng, et al., 2016). Such changes appear partially associated with poorer cognitive performance (Andrews-Hanna, et al., 2007; Chan, et al., 2014; Fjell, et al., 2015; Ng, et al., 2016; Salami, et al., 2014). However, the complete picture of whole-brain network activity and age-related changes in cognition across multiple domains is lacking.

The application of network measures to connectivity patterns has also revealed changes to the intrinsic functional architecture with age, namely a decreased modularity and segregation of RSN’s (Betzel, et al., 2014; Chan, et al., 2014; Geerligs, et al., 2015; Grady, et al., 2016). There also appears to be age-related decreases in connectivity for specific subnetworks of connections - particularly those involving long-range communication (Cao, et al., 2014; Marques, et al., 2015; Sala-Llonch, et al., 2014): These may reflect the decreased integration of large functional brain networks with age (Sala-Llonch, et al., 2014), consistent with the corresponding changes in structural networks (Perry, et al., 2015; Zhao, et al., 2015). Hitherto, only few investigations of intrinsic functional organisation in healthy elderly populations have been undertaken (Madhyastha and Grabowski, 2014; Marques, et al., 2015; Ng, et al., 2016; Sala-Llonch, et al., 2014). Although studies of age across the whole lifespan are illuminating and important, they typically contain relatively modest numbers of healthy older participants. Moreover, the association between cognitive performance and brain structural integrity is not uniform from adulthood to elderly years (Burzynska, et al., 2012; Razlighi, et al., 2016; Turner and Spreng, 2012). The later decades of life are also characterised by progressive changes in the performance of everyday functions (Ball, et al., 2007). There is hence a need to study the influence of age on functional connectivity patterns within a healthy elderly cohort.

Higher levels of educational attainment, intelligence, occupational status and other positive lifestyle factors contribute protective effects against age-related cognitive changes as well as the onset of AD (Deary, et al., 2009; Stern, 2012; Valenzuela and Sachdev, 2006). Expressions of these factors are postulated to contribute to an individual’s capacity to mitigate the influence of age, which has been broadly grouped together into the rubric term “cognitive reserve” (CR) (Stern, 2009; Stern, 2012). The proxies of CR are associated with a relative preservation of brain structure and more efficient neural activity during cognitive demands (Bartrés-Faz and Arenaza-Urquijo, 2011). Increased educational attainment is also associated with increased resting-state functional connectivity in distributed cortical networks (Marques, et al., 2016; Marques, et al., 2015). However, the influence of moderating factors such as education on the cognitive networks sensitive to age-related changes are poorly understood (Bartrés-Faz and Arenaza-Urquijo, 2011; Stern, 2009; Stern, et al., 2008). Some aspects of brain functions may be optimised with age (Moran, et al., 2014), which speaks to the potential adaptive influence of moderating factors on large-scale network interactions in later life.

Multivariate analyses allow a broad picture of brain-behaviour relationships. Using canonical correlation analysis (CCA), Smith and colleagues (Smith, et al., 2015) studied the complex inter-relationships between 158 phenotypic measures and whole-brain functional connectivity patterns in a large cohort of healthy younger adults (Van Essen, et al., 2013). Intriguingly, the co-variation between a full suite of phenotypic markers and functional connectivity loaded onto a single positive-negative axis. Positive personal traits (e.g. life satisfaction, education years, and fluid intelligence) shared strong co-variations with connectivity patterns, whilst characteristically negative traits (e.g. substance use) load negatively onto brain-behaviour associations. Related multivariate approaches such as partial least squares (PLS) (McIntosh, et al., 1996) have revealed latent patterns of functional activations related to decreased brain variability in older adults (Garrett, et al., 2011; Garrett, et al., 2012). Both CCA (Tsvetanov, et al., 2016) and PLS (Ferreira, et al., 2015) approaches have also recently revealed lifespan changes in functional connectivity patterns. These findings are derived from cohorts that span the whole lifespan and did not address the relative influence of age on neurocognitive networks. Some cognitive functions – such as psychomotor abilities – are more susceptible to age-related changes in later life than others (Salthouse, 1996). The influence of age is likely to act most strongly on networks supporting these functions.

Multivariate techniques have not yet been employed to investigate these issues, nor the relative influence of both age and CR proxies on connectivity patterns. Here we use multivariate analysis to examine the associations between age, education, cognitive performance (measured across a number of domains) and whole-brain functional connectivity patterns in 101 cognitively-normal elders. In particular, we ask whether the single positive-negative axis of associations between behavioural indicators of cognition and functional brain networks seen in young adults (Smith, et al., 2015), persists under the influence of healthy ageing. We hypothesise that connectivity patterns associated with cognitive domains most susceptible to decline such as processing speed will most be strongly opposed to the influence of age. We also ask whether moderating factors such as education confer an influence upon age-varying networks, or rather onto independent brain-behaviour modes.

## Materials and methods

### Participants

Cognitively-normal individuals were drawn from a longitudinal, population-based study (the Sydney Memory and Ageing Study (MAS) (Sachdev, et al., 2010a)). At the baseline of this longitudinal study, community-dwelling participants initially between 70-90 years of age were randomly recruited from the electoral roll. Imaging and phenotypic data for the present paper were acquired during the fourth wave of this study (approximately 6 years following study baseline). Subjects were classified as cognitively normal at the current wave if their performance on all neuropsychological test measures was higher than a threshold of 1.5 SDs below normative values. The criteria for the selection of this cohort, and the demographic matching that was used to establish a normative reference have been previously published (Tsang, et al., 2013). Exclusion criteria for all study participants at baseline included a Mini-Mental Statement Examination (MMSE) (Folstein, et al., 1975) adjusted (Anderson, et al., 2007) score below 24, a diagnosis of dementia, developmental disability, a history of schizophrenia, bipolar disorder, multiple sclerosis or motor neuron disease, active malignancy, or inadequate comprehension of English to complete a basic assessment. 135 participants with concurrent MRI data met inclusion criteria. The study was approved by the Ethics Committee of the University of New South Wales and participants gave written, informed consent.

### Neuropsychological measures

A comprehensive neuropsychological battery was administered by trained graduate psychologists to cover a broad range of cognitive functions, including attention, processing speed, memory, language, visuospatial ability, and executive function. Twelve tests were grouped into five broad domains - attention/processing speed, memory, language, visuospatial ability, and executive function (Table 1). Each domain consisted of a composite of these individual tests, with the exception of the visuospatial domain which was represented by a single measure. As part of the broader longitudinal study (MAS) (Kochan, et al., 2010; Sachdev, et al., 2010b) - the tests were grouped according to the primary cognitive function they assess - based upon the extant literature and the widespread practice used by neuropsychologists (Lezak, et al., 2004; Strauss, et al., 2006; Weintraub, et al., 2009). The groupings align with the domains of established psychometric batteries such as the UDS (ADC) (Morris, et al., 2006; Weintraub, et al., 2009), and other large epidemiologic cognitive ageing studies (Mayo study (Roberts, et al., 2008); MYHAT study (Ganguli, et al., 2010)). The memory domain composite was further subdivided into verbal memory after exclusion of a visual retention test (Benton, et al., 1996). We additionally study the relative weighting of each individual neuropsychological test onto our primary results.

**Table I.** Neuropsychological tests administered to measure the cognitive grouping scores

| **Neuropsychological Test** | **Cognitive Grouping** |
| --- | --- |
| • Digit Symbol-Coding (Wechsler, 1997a)<br>• Trail Making Test (TMT) A (Strauss, et al., 2006) | **Attention/Processing Speed** |
| • Logical Memory Story A delayed recall (Wechsler, 1997b)<br>• Rey Auditory Verbal Learning Test (RAVLT) (Strauss, et al., 2006):<br>○ RAVLT total learning; sum of trials 1-5<br>○ RAVLT short-term delayed recall; trial 6<br>○ RAVLT long-term delayed recall; trial 7<br>• Benton Visual Retention Test recognition (Benton, et al., 1996) | **Memory** |
| • As above, but not including the Benton Visual Retention Test. | **Verbal Memory** |
| • Boston Naming Test – 30 items (Kaplan, 2001)<br>• Semantic Fluency (Animals) (Strauss, et al., 2006) | **Language** |
| • Block Design (Wechsler, 1981) | **Visuospatial Ability** |
| • Controlled Oral Word Association Test (Strauss, et al., 2006)<br>• TMT B (Strauss, et al., 2006) | **Executive Function** |

To further support the cognitive groupings, we performed reliability estimates which measure the scale-item’s homogeneity (SI Table 1). For the full healthy reference cohort (n = 343; with no missing domain scores), reliability estimates reveal acceptable (*r_SB_* > 0.70) to high internal consistency of the composite-items - according to psychometric convention (Cortina, 1993; Tavakol and Dennick, 2011). The only exception was the executive function composite. Tasks of executive functions (as with the other domains) posses a multifactorial structure as they rely on other cognitive systems/processes for their expression (Piguet, et al., 2005), and hence the lower item-homogeneity here is not surprising. For example, the Trail Making Task (TMT) B is also associated with lower-order abilities (Sanchez-Cubillo, et al., 2009). Performance on the other executive composite, the FAS, also relates to verbal intelligence, lexical retrieval and processing speed (Greenaway, et al., 2009).

The individual test scores for each subject were transformed into quasi *Z*-scores based upon the mean and standard deviation of tests scores for a healthy, reference group (*n* = 723) phenotyped at study baseline. Domain scores were calculated as the average of the quasi *Z*-scores of tests comprising each domain. If necessary, the signs of the z-scores were reversed so that higher scores reflect better performance.

Clinical measures including the MMSE and the Bayer-Activities of Daily Living Scale (B-ADL) (Erzigkeit, et al., 2001; Hindmarch, et al., 1998) were also administered. The B-ADL consists of 25 informant-rated items - scored on a scale of 1-10 - according to the frequency of participant’s impairments in everyday activities. Higher scores on the B-ADL relate to more severe deficits in functioning. The mean values for each participant were defined by the average score across the B-ADL questionnaire items. These clinical ratings were used in the present study for further characterization of the current samples cognitive and functional status. The B-ADL scores for 17 study participants were however missing.

The National Adult Reading Test (NART IQ) (Nelson and Willison, 1991) was administered to a subset of the current population at study baseline. The NART estimates premorbid intelligence levels (Bright, et al., 2002).

### Acquisition and pre-processing of MRI data

Eyes-closed resting-state fMRI (rs-fMRI) data consisting of 208 time-points were acquired with a T2* weighted echo-planar imaging sequence (TE = 30 ms, TR = 2000 ms, 1.87 x 1.87 x 4.50 mm^3^ voxels) on a Philips 3T Achieva Quasar Dual MRI scanner (Amsterdam, the Netherlands). Structural T1-weighted MRI were also acquired (TR = 6.39 ms, TE = 2.9 ms, 1mm^3^ isotropic voxels). FSLView (Smith, et al., 2004) was used to visualise every MRI scan for artifact inspection. Subjects were removed if their data contained excessive artefact, including the presence of complete orbitofrontal signal dropout (Weiskopf, et al., 2007), motion effects on T1-images (i.e. ringing), or severe geometric warping. A full description of the steps involved for the acquisition and pre-processing of these data are provided in Supplementary Information (SI) 1.1 and 1.2.

Data pre-processing was performed using the Data Processing Assistant for Resting-State fMRI (DPARSF, v3.2 advanced edition) software package (Yan and Zang, 2010), which calls functions from SPM8 (http://www.fil.ion.ucl.ac.uk/spm/). Basic pre-processing steps included slice-timing, realignment to mean functional image, co-registration of the structural image, linear detrending, and nuisance regression of head motion (24 parameters) (Friston, et al., 1996) and segmented WM/CSF signals (Ashburner and Friston, 2005). Native functional images were transformed into an average population-based T1 template (i.e. DARTEL) (Ashburner, 2007) and then Montreal Neurological Institute (MNI) space (3mm^3^ voxels). fMRI images were smoothed (at 8mm) and temporal band-pass filtering applied (0.01– 0.08 Hz). Global signal regression was not performed.

Of the initial subject population (*n* = 135), fifteen were removed due to severe signal loss (thirteen within fMRI scans), ten had incomplete cognitive information, whilst nine failed adequate co-registration between their T1-weighted and mean functional image. Data from one-hundred and one subjects were hence included in the primary analysis (Table II).

**Table II.** Basic demographic, cognitive and clinical information for included participants

| Cohort | All subjects ($n = 101$) |
| --- | --- |
| NESB ($n$) | 10 |
| M/F ($n$) | 44/57 |
|  | Mean (+- SD) |
| Age (years) | 82.7 (3.8) |
| Education (years) | 12.7 (3.6) |
| MMSE | 29.5 (0.9) |
| B-ADL* | 1.41 (0.54) |

NESB, Non-English Speaking Background; M, Male; F, Female; B-ADL, Bayer-Activities of Daily Living Scale
\* $n = 19$ participants were missing data

### Construction of functional brain networks

In brief, the Pearson’s correlation coefficient of the mean BOLD signals between all pairs of 512 uniformly-sized regions (SI 2) (Perry, et al., 2015; Zalesky, et al., 2010) was calculated to construct the functional connectivity matrix *M.* Fisher’s transformation was applied to *M*, and subsequent upper-triangle values were concatenated across all subjects, forming a matrix *N*_1._ Full description of the steps involved for the construction and normalization of functional brain networks are provided in SI 1.3 and 1.4.

### Normalization, demeanining and head-motion regression of connectivity matrices

The connectivity matrices *N*_1_ were normalized and demeaned according to the procedure of (Smith, et al., 2015) (also available online at http://fsl.fmrib.ox.ac.uk/analysis/HCP-CCA/hcp_cca.m), resulting in a matrix *N*_2_ for subsequent analyses. The mean frame-wise displacement (FD) (Power, et al., 2012) was calculated and potential confounding effects of head motion were regressed from *N*_2_ to yield *N*_3_. Notably, there was no significant relationship between age and FD (*p*>0.39, *r* = -0.09).

### Functional connectivity decomposition

Principal Components Analysis (PCA) was implemented via the FSLNets toolbox (Smith, et al., 2014) to reduce the dimensionality of the functional connectivity edges (*N*_3_) to eight eigenvectors. Given that eight non-imaging measures were selected in the CCA (see below), the data was reduced to this resolution to keep the methodological steps similar to Smith et al. (2015); In their study, the greatest fit (correlation) between the connectivity and non-imaging weights was obtained by the CCA which used the same number of brain and phenotypic components. However, no gold standard exists for component number selection (Abdi and Williams, 2010). We note that the first eight functional components explain 20.3% of the total proportion of variance (SI Fig 1, red bars).

### Canonical Correlation Analysis (CCA)

Eight subject measures were chosen for inclusion in the CCA: Age, education years, and the composite scores for language, executive function, visuo-spatial ability, memory, verbal memory and attention/processing speed. NART IQ scores were administered only to a subset of the current cohort (*n* = 91) at wave 1.

CCA was then applied to these non-imaging measures and functional eigenvectors, yielding eight modes which constitute weighted linear combinations of orthogonalized subject measures and functional connectivity patterns: Each mode represents canonical correlations which correspond to the maximum residual co-variation between the two variate sets in decreasing rank order. The vectors *U_m_* and *V_m_*, represent the individual subject weights for subject measures and connectivity matrices within mode *m* respectively:

- *U_m_* is the extent to which each subject is (positively or negatively) correlated to population variation in subject measures within mode *m*
- *V_m_* is the extent to which each subject is correlated to population variation in brain connectivity within mode *m*

The correlation of *U_m_* and *V_m_* yields *r_m_*, the strength of the population co-variation in mode *m* shared between brain connectivity and subject measures.

To assess the reliability of the loading of cognitive and demographic measures onto each mode *m*, a bootstrapping procedure (sampling with replacement) was performed over 5000 subsamples. The phenotypic loadings within each mode *m* were considered reliable if the 95% confidence bounds of the bootstrapped distribution of correlations did not overlap with zero (Ferreira, et al., 2015; McIntosh, et al., 2004).

### Association of connectivity edges within each mode

We next assessed which connectivity edges were most strongly expressed by population variations in connectivity captured within mode *m*. First, to obtain the relative weight (and directional signs) of each edges association with the connectivity patterns within mode *m*, we correlated *V_m_* with the original connectivity estimates in *N*_3_, resulting in a vector *A_Fm_*. The connectivity edges identified most strongly associated with either positive or negative co-variations between *U_m_* and *V_m_*, were chosen as the top 250 strongest connections (representing 0.002% of all network edges) with positive and negative signs within *A_Fm_* respectively.

### Publicly-available code

The MatLab codes implemented for the normalization and PCA of the connectivity edges - as well as the steps involved in the CCA - are stored in a publicly-available repository (https://github.com/AlistairPerry/CCA). The repository additionally contains further information for the brain parcellation templates used in functional network construction.

### Statistical Analyses

To determine the significance of interdependence between the variates sets within each mode *m*, Wilk’s Lambda was first calculated and transformed into Rao’s Approximation *F*-statistic (Rao, 1952). Shared variances captured between the respective variate sets of mode *m* were determined as significant if *p*<0.05, thus rejecting the null hypothesis (H0) that subject measures and functional components are independent of each other within mode *m*.

## Results

Our cohort of 101 cognitively-normal healthy elders span the later decades of life. Raw performance on the neuropsychological tests and cognitive grouping scores are provided in SI Table II and SI Table III, respectively. The clinical rating scores of the current population are within the range of values for previously published data of healthy older adults (Table II) (Erzigkeit, et al., 2001; Reppermund, et al., 2011; Roalf, et al., 2013). A clear association between age and cognitive performance is demonstrated, particularly for attention/processing speed scores (Fig 1A, top-left panel; *p*<0.001, FDR-corrected). In the full sample, we also assessed the complex relationships between age, sex (males coded as 1), education and six cognitive groupings: verbal memory, memory, visuospatial ability, executive function, language and attention/processing speed. Performance across these cognitive groupings is highly correlated (Fig 1B). Performance in visuospatial, executive function and language domains is positively correlated with years of education (*p*<0.05; FDR-corrected). As expected, memory and verbal memory (being largely redundant) correlate very strongly. Memory performance is significantly correlated with female sex (Fig 1B; *p*<0.001, FDR-corrected), whilst males demonstrate greater visuospatial abilities (*p*<0.05, FDR-corrected). For the subset who received NART IQ assessment at study baseline, we also examined relations with IQ (SI Fig 2). There are no significant differences (*p*<0.05, two-tailed, FDR-corrected) between the full study cohort and this subset population across the phenotypic variables (SI Table III).

**Fig 1.**
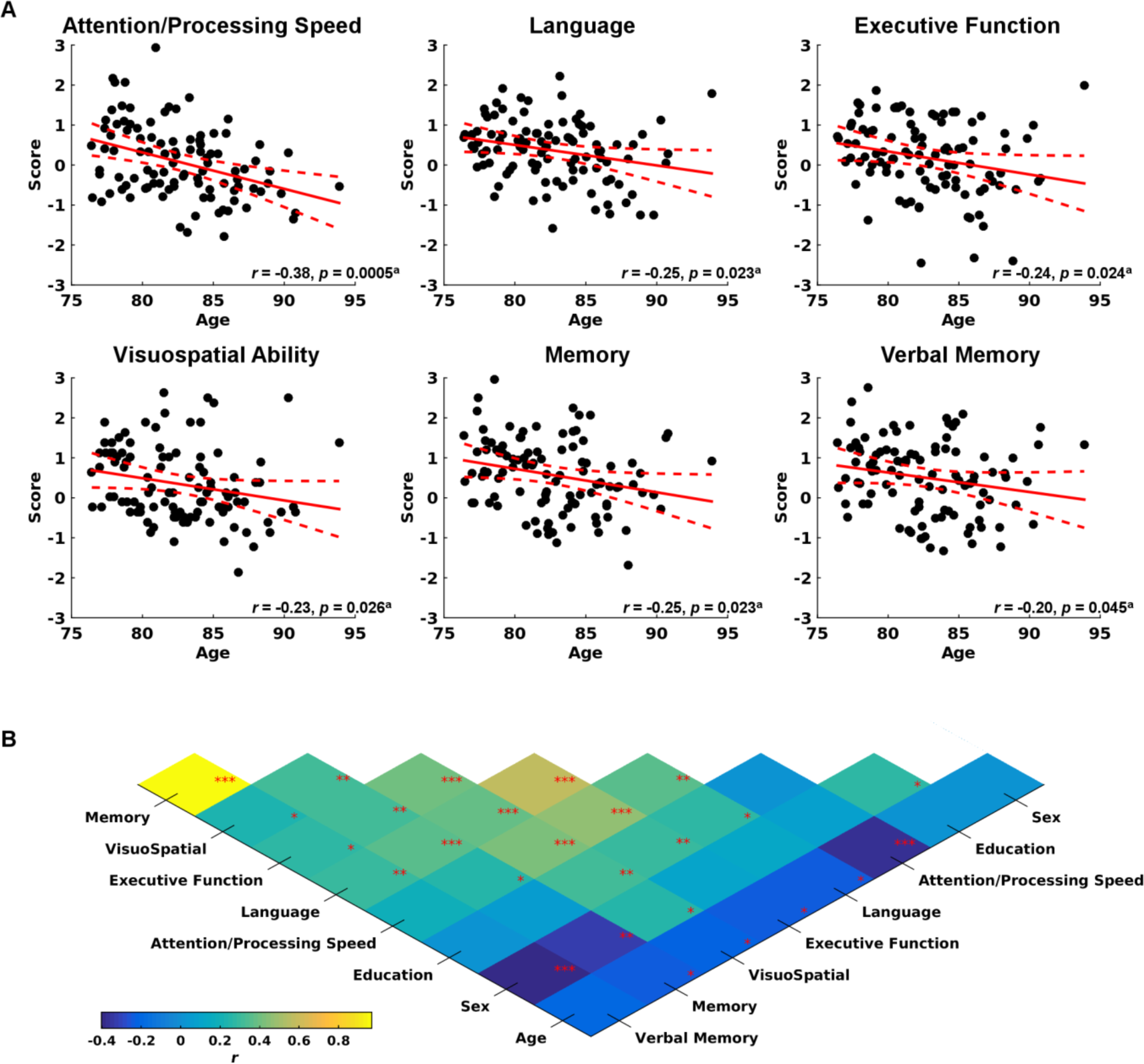
Associations between the phenotypic information of the healthy older adults. (A) Cognitive functioning across the groupings as a function of age. Solid red lines show the best-fitting linear regression of age, whilst dashed red lines indicate the 95% confidence intervals for the linear fit. (B) Strength and direction of associations between all phenotypic measures. *a*, FDR-corrected * *p* < 0.05, ** *p* < 0.01, *** *p* < 0.001; FDR-corrected N.B: Males coded as 1

We next used CCA to examine the primary modes that relate these (correlated) demographic and cognitive variables to functional connectivity patterns. CCA identified three significant canonical modes (*p*<0.05) of interdependence between these non-imaging measures and functional connectivity (Table III).

**Table III.** Significant (*p*<0.05) CCA modes in the primary analysis

| CCA Mode | One | Two | Three |
| --- | --- | --- | --- |
| $df_1$ | 64 | 49 | 36 |
| $df_2$ | 496.76 | 441.03 | 384.80 |
| $F$ | 1.77 | 1.55 | 1.48 |
| $\lambda$ | 0.30 | 0.45 | 0.57 |
| $r^2$ | 0.32 | 0.21 | 0.20 |
| RI | 0.072 | 0.030 | 0.023 |
| $p$ | 0.00043 | 0.012 | 0.041 |

$\lambda$ = Wilk's lambda; RI = redundancy

Each CCA mode consists of a set of weights that reflect the loading of the cognitive and demographic variables onto the corresponding functional connectivity patterns (Fig 2). The first mode (*p*<0.00043) is characterised by a split between all cognitive domains (particularly memory and attention/processing speed) which load along a positive axis, and age which loads strongly and negatively (Fig 2A, left panel). Language and education have close to zero loading and are not reliably represented within this mode (i.e. the confidence bounds of the bootstrapped distributions cross zero; Fig 2A, grey text). The opposing pull of attention and processing speed versus age can be seen by plotting the subject specific measure weights versus the corresponding connectivity weights, coloured according to age (Fig 2D) or attention/processing speed (Fig 2E). Younger subjects (Fig 2D, blue circles) cluster in the top right corner of the panel, indicating how they weigh positively with the corresponding connectivity-behaviour relations. Likewise, fast and attentive performers (Fig 2E, green to dark red) also load positively on the first CCA mode. These plots show that faster, attentive, younger performers weight positively with functional connectivity patterns within this mode, whereas poorer, older performers contribute to negative associations.

**Fig 2.**
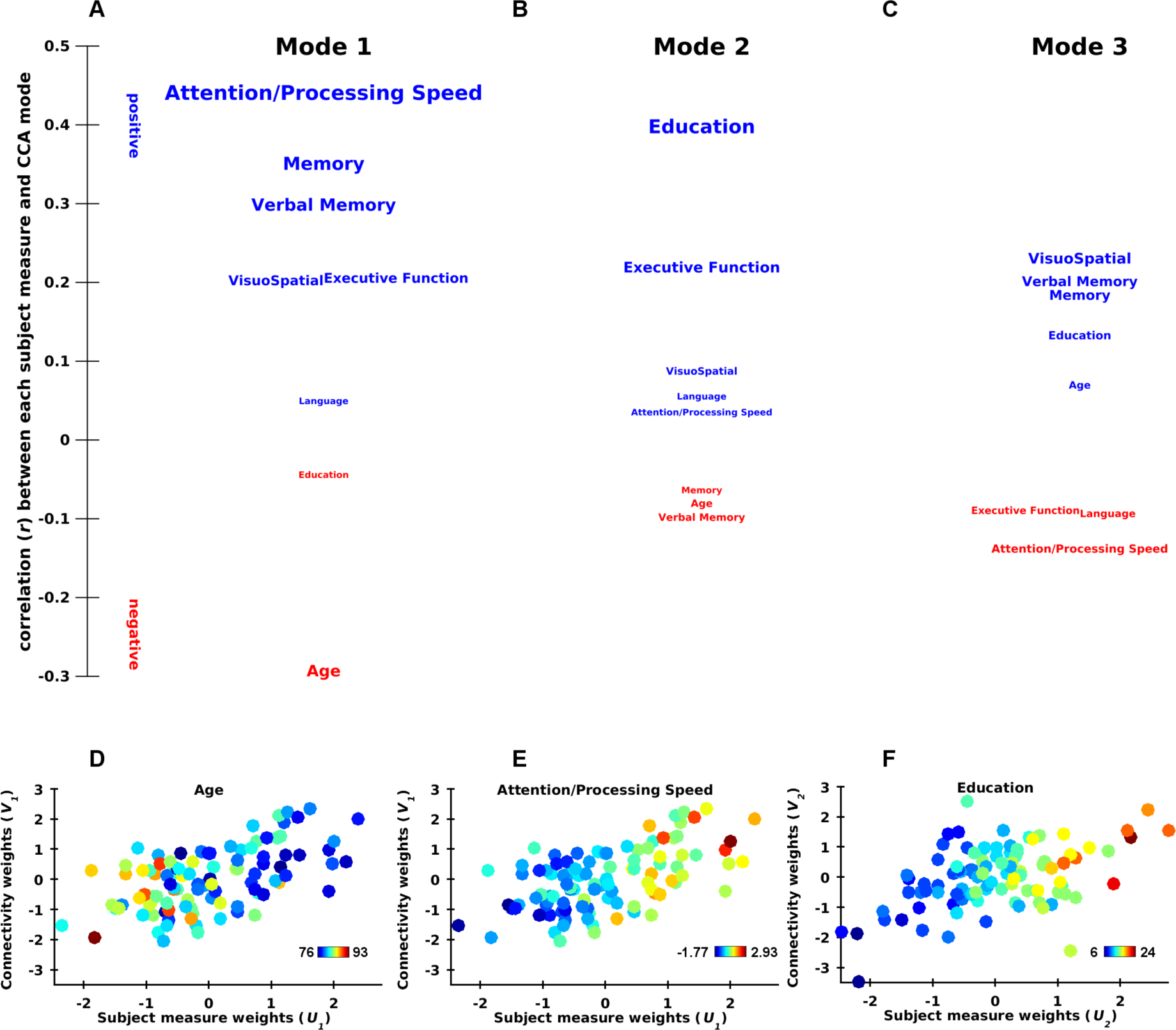
Weighting of cognitive and demographic measures captured by the CCA modes. (A-C) Correlation between subject measures and functional connectivity captured by the mode (*V_m_*), with the strength and direction indicated by the vertical position and font size. Grey text depicts phenotypic loadings where the confidence intervals of bootstrapped distributions overlap with zero. (D-F) Scatter plots showing for each subject (data points) their weighting towards non-imaging measures (*U_m_, x*-axis) and functional connectivity patterns ( *V_m_, y*-axis), captured for the first (D-E) and second modes (F). Colour is scaled according to subjects age (D), Attention/Processing Speed performance (E), and education level (F).

In contrast, the second mode (*p*<0.012) is characterised by an independent positive association of education years with connectivity patterns (Fig 2B, 2F). Although executive function loads moderately on this mode, all other variables load very weakly (in both directions). While age and memory load negatively, their contributions are weak. There also exists a weakly significant third mode (*p*<0.041). This mode splits cognitive measures into moderately positive visuospatial and memory weights versus weakly negative attention and processing speed (Fig 2C). Age and education weigh close to zero.

Each of the three CCA modes also load onto patterns of functional brain connectivity. To study these, we calculated the 250 edges most strongly associated with each mode in both the positive and negative directions. The functional connectivity edges most strongly expressed by positive associations in the first mode (mean *r* = 0.64, *SD* = 0.02) primarily involve bilateral connections between occipital, temporal (inferior and medial portions), superior parietal, and pre/post central gyral regions (Fig 3A). Functional connections between occipital areas and pre/post-central regions within the right hemisphere are also evident. Of note is the convergence of connections upon bi-lateral parietal operculum/posterior insular areas. To disentangle the functional basis of this network of strongly associated connections, we assigned regions in our parcellation to broader functional network clusters; default-mode, cognitive-control, somatomotor, dorsal attention, salience ventral attention, visual, and limbic networks (Yeo, et al., 2011). This demonstrates positive edges in the first mode are predominately clustered among visual, somatomotor, and to a lesser extent, dorsal attention networks (Fig 3A and 3B).

**Fig 3.**
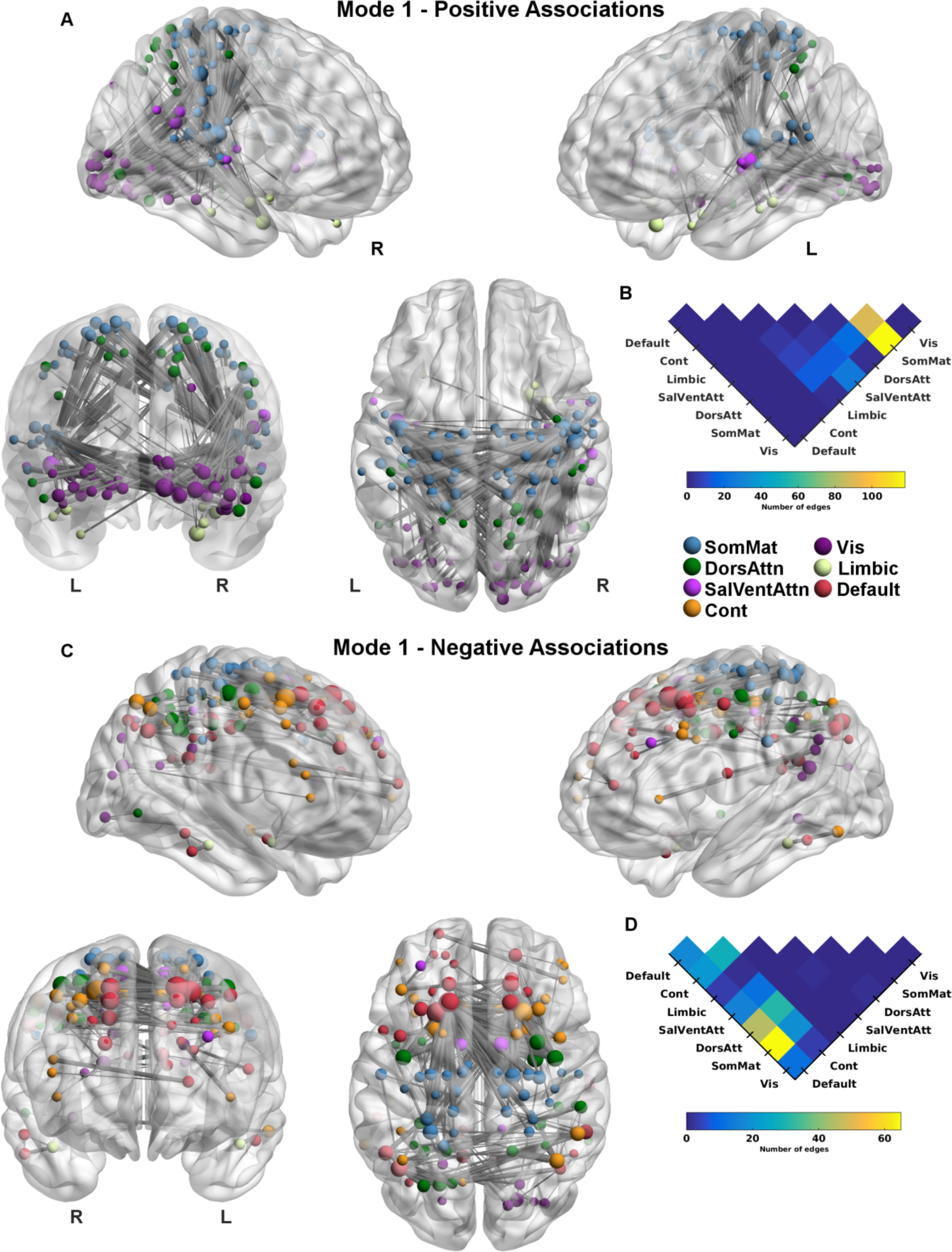
Connectivity edges most positively and negatively expressed by the first CCA mode. (A) and (C) Connectivity edges exhibiting the strongest positive and negative associations with functional connectivity patterns (*V_1_*), respectively. Line width indexes strength of correlation. Circle size is scaled to the number of connections each region shares within the network; Node colour denotes functional network affiliation (Yeo, et al., 2011). The brain meshes are presented from axial (bottom middle panel), coronal (bottom left), and customised perspectives of the left (top right; elevation = 0, azimuth = -120) and right-hemisphere (top left; azimuth = -240). Connectivity edges and surface meshes were visualised using BrainNet Viewer (Xia, et al., 2013). (B) and (D) Connectivity distributions across the functional clusters for the edges most positively and negatively expressed. Warmer colours indicate greater number of connections

We then identified the functional connectivity edges most negatively expressed by the first mode (*mean* = -0.27, *SD* = 0.03). These connections form two distinct clusters: The first cluster inter-connects pre-motor, pre/post central gyri and superior medial frontal areas (supplementary motor area, pre-supplementary, superior frontal gyri) (Fig 3C). A second cluster involves inter-hemispheric connections between inferior parietal areas, and additional connections between these areas and left superior parietal regions. On a coarser scale these edges connect DMN and cognitive-control network areas to regions affiliated with all other networks except for limbic areas, particularly default-mode connectivity with both the somatomotor and dorsal attention networks (Fig 3D).

The edges most strongly expressed within the second mode are quite distinct from those demonstrating positive associations with the first mode, mirroring the divergent loading of phenotypic measures. The edges exhibiting the strongest positive associations (*mean* = 0.73, *SD* = 0.01) with the second mode stretch between visual cortices and dorsolateral prefrontal areas, whilst connections from superior parietal (dorsal attention) and pre/post-central gyri (somatomotor) converge at both dorsolateral and ventrolateral regions, within default and control networks (Fig 4A). Assigning regions to their respective functional affiliation shows that edges from the default and control networks inter-connect preferentially with visual, somatomotor, and dorsal attention networks (Fig 4B).

**Fig 4.**
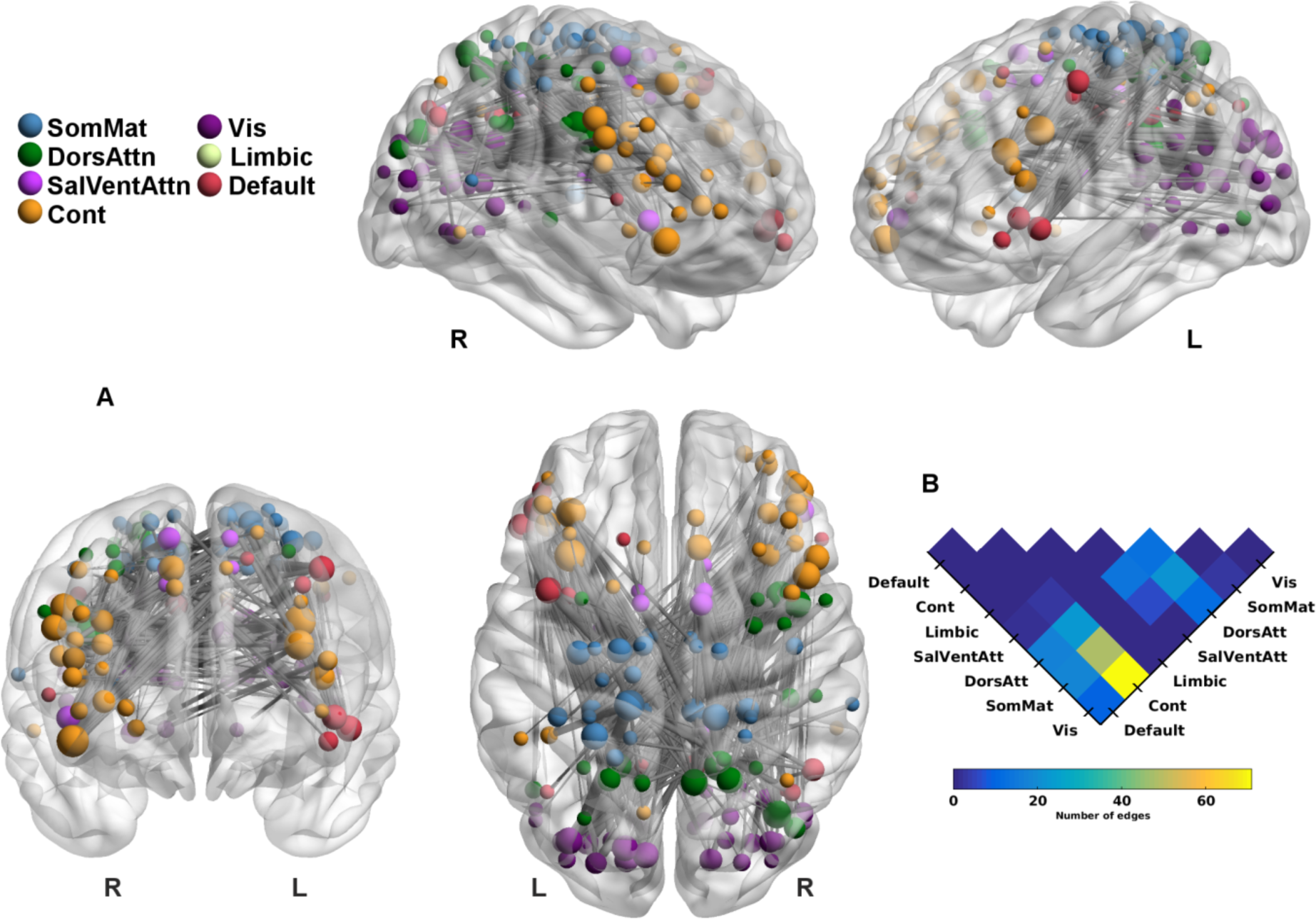
Connectivity edges most positively expressed by the second CCA mode. (A) Connectivity edges exhibiting strongest positive associations with functional connectivity patterns (*V_2_*), hence representing connections expressed by the increased education level. Line width indexes strength of correlation. Circle size is scaled to the number of connections each region shares within the network, whilst coloured to their functional network affiliation. The brain meshes are presented from axial (bottom middle panel), posterior (bottom left), and angular perspectives of the left (top right) and right-hemisphere (top-left). (B) Connectivity distribution across the functional networks, with warmer colours indicating greater number of connections.

The edges exhibiting the strongest positive associations (*mean* = 0.64, *SD* = 0.02) with the third mode also comprise networks that are distinct from the other two modes. Functional connections predominately cluster around ventrolateral and orbitofrontal divisions of left prefrontal nodes encompassing default-mode, cognitive control, and limbic areas (SI Fig 4A). Edges stretch between these areas and bi-lateral frontomedial regions (anterior cingulate and superior portions), the left cingulate (middle and posterior portions), and left inferior parietal lobe. Assigning these networks to functional subdivisions of the brain shows they are predominately distributed within-and-between default-mode and control network areas, with additional connections between all other networks (except for visual) (SI Fig 4B).

### The influence of sex and intelligence

Given the strong correlations between sex and cognitive performance across specific domains (Fig 1), we undertook an additional CCA with sex (males coded as 1) included (hence now with nine functional components). Two significant CCA modes were identified (*p* < 0.05, SI Table IV), showing subtle differences to the principal modes explored above (SI Fig 5). In the first mode (SI Fig 5A), cognitive domains are again spread along the positive axis, with (male) sex loading most strongly on the negative axis followed by age and education years. The strong independent association of education with connectivity remains in the second mode (SI Fig 5B), where sex and the cognitive domains demonstrate weak to moderate associations. The functional connections most strongly expressed by the first mode when including sex in the CCA are spatially consistent with those identified within the primary analysis (Fig 5A-B; red lines indicate edges that are strongly expressed regardless of whether sex is included in, or excluded from the CCA model). Analysis of the second mode, however, reveals edges that are predominantly expressed only when including sex within the model (Fig 5C; grey lines).

**Fig 5.**
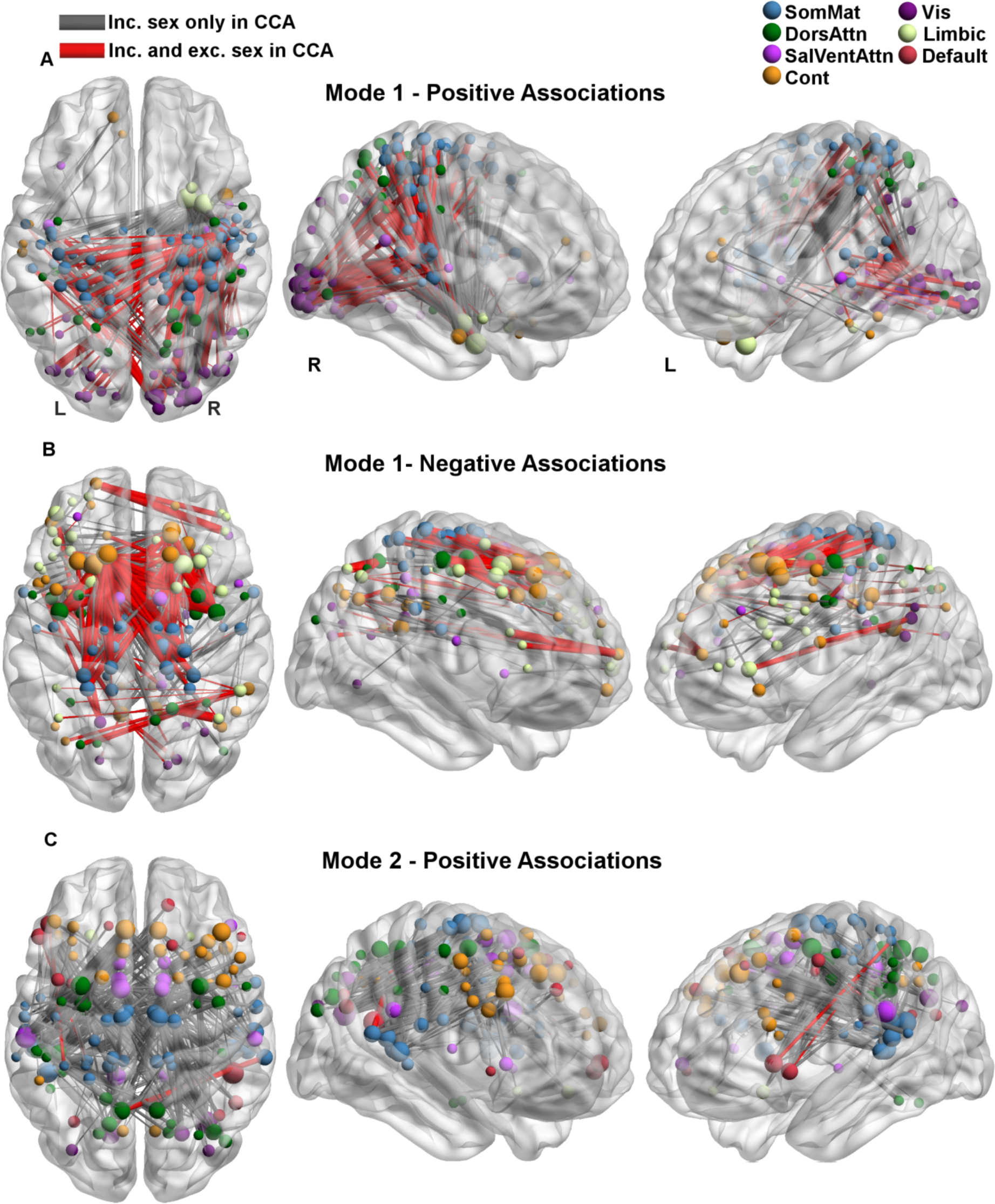
Connectivity edges most strongly expressed by the significant modes when including sex in the CCA model. (A) and (B) Connectivity edges exhibiting the strongest positive and negative associations with the functional connectivity patterns of the first mode (*V_1_*), respectively. (C) Connectivity edges exhibiting the strongest positive associations with the second mode (*V_2_*). Red lines indicate edges which are strongly expressed by CCA models with and without sex included, whilst grey lines are those uniquely expressed by the CCA with sex included. Line width indexes strength of correlation. Node size is scaled to the number of connections each region shares within the network; Node colour denotes their functional network affiliation. The images are presented from axial (left panel), and angular perspectives of the left (right) and right-hemisphere (middle).

Education and intelligence (as estimated by NART IQ scores) are highly-correlated (SI Fig 2), and both considered central to cognitive reserve (Stern, 2009). The positive co-variation between years of education and increased connectivity captured by the second mode in the main analyses thus raises an interesting question regarding the contribution of intelligence. We thus performed CCA (again with nine functional components) using the full cohort of subjects whom received NART IQ assessment at study baseline (*n =* 91). This analysis yielded two significant modes (*p*<0.05; SI Table V). The modes capture latent relations that are similar to the primary analysis, although interesting differences between education and intelligence emerge (Fig 6A-B). Within the first mode, NART IQ loads positively and of similar magnitude to memory and visuospatial ability. Although NART IQ scores also bear a moderate positive association with connectivity captured by the second mode, the strength of this loading is weaker than education. Thus NART IQ splits across modes, with some weighting in opposition to age and some loading independently with education.

**Fig 6.**
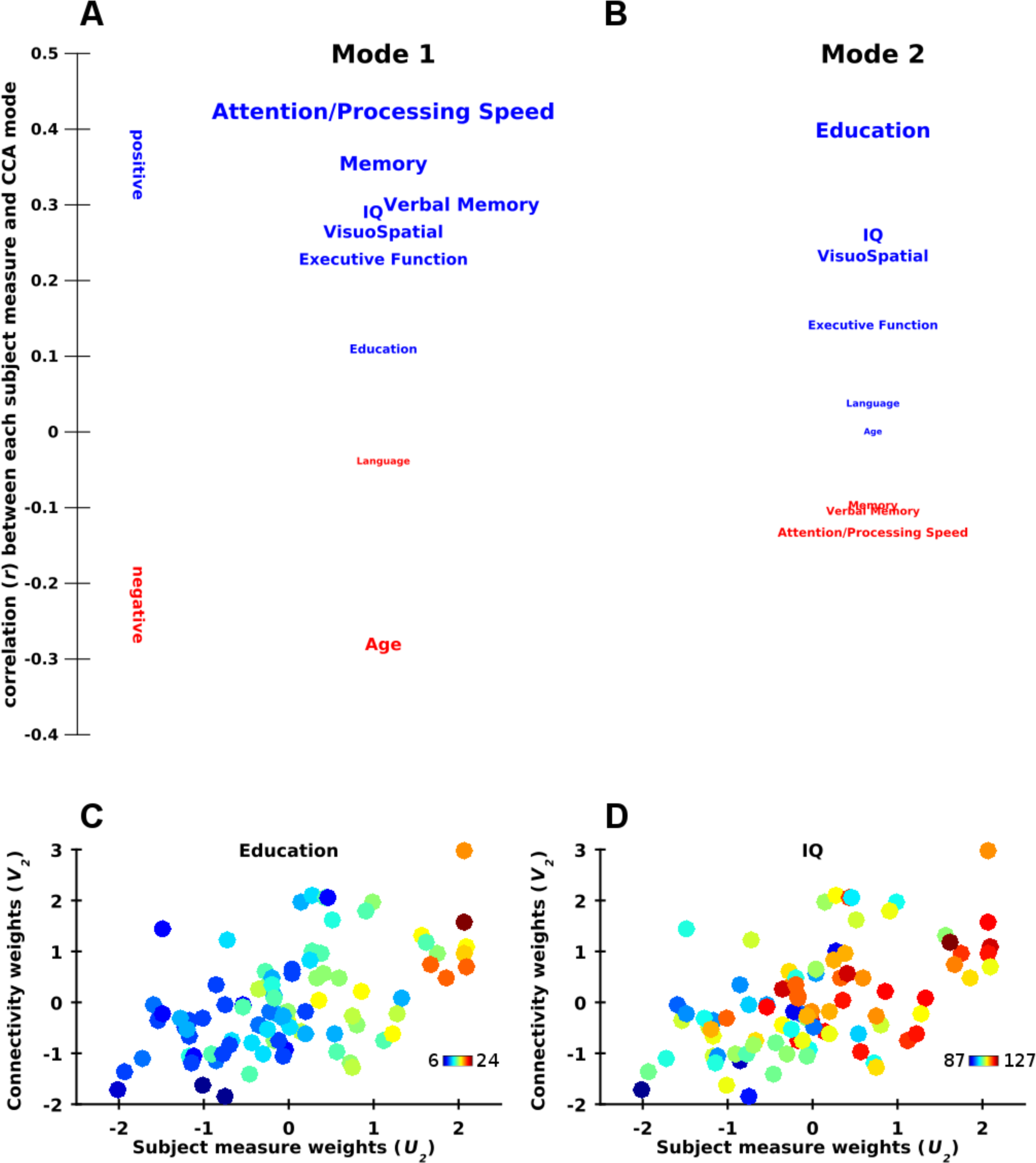
Weighting of cognitive and demographic measures captured by the CCA modes including intelligence scores. (A-B) Correlation between subject measures and functional connectivity variation (V_m_), with the strength and direction indicated by vertical position and font size. (C-D) Scatter plots showing for each subject (data points) their weighting towards non-imaging measures (*U*_2_, x-axis) and functional connectivity patterns (V_2_, y-axis), captured for the second modes. Colour is scaled according to subject’s education level (C) and NART IQ scores (D).

The strong influence of education when also including IQ in the CCA model, also raises an interesting question regarding the functional connectivity patterns that are captured here. The edges exhibiting the strongest positive associations (*mean* = 0.74, *SD* = 0.015) are distributed throughout the cortex (Fig 7). Several key features are evident: Connections converge (larger circles) upon parietal default-mode areas including right -and medial (precuneus) portions, as well as superior (dorsal attention), and paracentral areas (somatomotor). Edges connect these areas to DMN and dorso- and ventrolateral prefrontal areas as well as lateral pre- and postcentral gyri. Only a small proportion of function connections (24/250 edges = 9.6%) also occur within the corresponding mode of the primary analysis (SI Fig 12). Visualising this network with a connectivity heat map (Fig 7B) and edge bundling connectogram (Fig 7C) (https://cran.rproject.org/web/packages/edgebundleR/index.html) which acts to cluster hierarchical relationships, shows that edges predominately cluster between default-mode (red circles) and control-network (orange) areas to all other networks except for limbic regions. Notably, the edges cluster around key DMN and control-network regions (larger circles).

**Fig 7.**
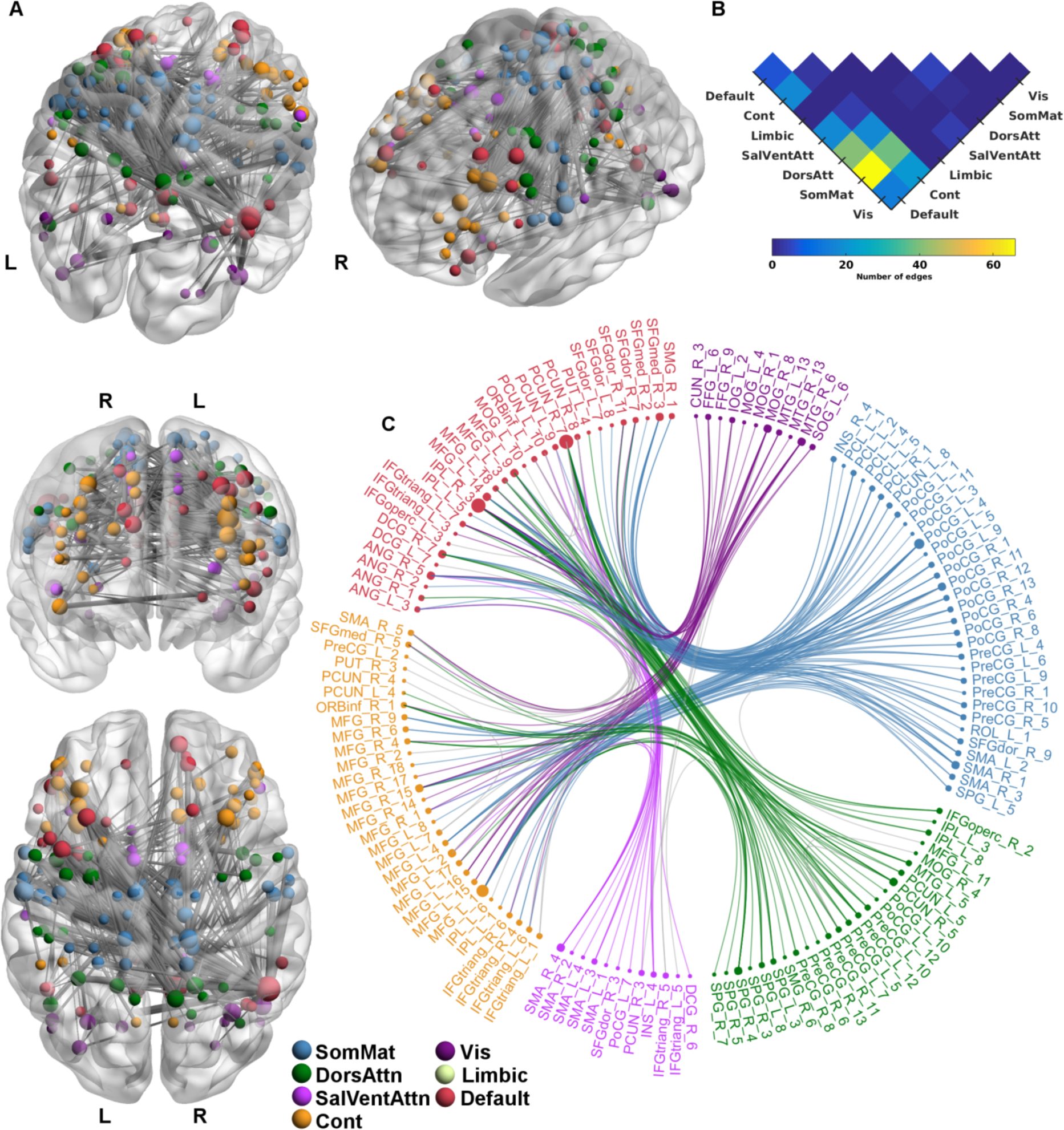
Connectivity edges most positively expressed by the second CCA mode including intelligence scores. (A) Connectivity edges exhibiting strongest positive associations with functional connectivity patterns (*V_2_*). Line width indexes strength of correlation. The brain meshes are presented from axial (bottom-left panel), posterior (middle-left), and customised superior (top-middle; elevation = 55, azimuth = 18) and lateral (top-right; elevation = 40, azimuth = -100) perspectives. (B) Connectivity distribution across the functional clusters, with warmer colours indicating greater number of connections. (C) Edge-bundling connectogram which clusters the hierachial relationships between these set of connections. Positions of regions are according to their network affiliation. Edges are coloured by their respective affiliation if they link to either default-mode or control-network regions. All other possible interactions are coloured grey. For both (A) and (C), circle size is scaled to the number of connections each region shares within the network; Circle is coloured to their functional network affiliation.

### Auxiliary analyses: Removing verbal memory, head motion confounds, functional eigenvectors, parcellation scheme, smoothing kernel

The construct of memory in the primary analysis includes verbal memory and is hence partly redundant (and thus strongly correlated) when verbal memory is also coded separately. However, two significant CCA modes (SI Table VI) were also identified with the removal of verbal memory scores with almost identical loading distributions to those in the original analysis (SI Fig 6).

The potential confounds of head motion were already regressed from the analysis. Nonetheless, further validations were also performed. Subject motion (i.e. mean FD) shows no significant association with functional connectivity variation (i.e. *V_m_*) captured across all three modes (*p*>0.95, FDR-corrected; SI Fig 7A-C), nor does it influence brain-behaviour relations (SI Fig 7D-F).

To determine that the captured brain-behaviour relations are not dependant on the amount of functional connectivity information which is fed into the CCA, we re-performed the analysis with four functional components; A shoulder in the variance contribution is apparent close to the fourth eigenvector (SI Fig 1) – which according to the subjective scree/elbow test (Abdi and Williams, 2010; Cattell, 1966) - represents sufficient variation captured from the connectivity data: Two significant CCA modes were identified (SI Table VII), with almost identical loadings (SI Fig 8) to the primary analysis. The additional significant mode when using the eight modes likely captures some residual covariance adjusting the main two modes to the residual variance.

To check the dependence of our findings on the parcellation scheme employed, the analysis was repeated, using a coarser brain template of 200 regions (including cerebellar and brain stem areas) derived by spatial-clustering of functional connectivity patterns in an independent data set (Craddock, et al., 2012). The positive-negative split of cognitive domains and age remains present within the first modes albeit the significance is slightly reduced (SI Table IX, SI Fig 9A). The independent loading of education on the second mode also remained, although this was again slightly reduced (SI Fig 9B). Visual inspection of the connectivity edges that are most strongly expressed with implementation of a coarser template reveals a spatial distribution that is consistent to that identified within the fine-grained parcellation (SI Fig 10). Two significant modes were also identified when a 6mm smoothing kernel was applied to our rs-fMRI data (SI Table X, SI Fig 11).

## Discussion

We used a multivariate approach to reveal the complex relationships between demographic factors, cognitive performance and functional brain networks in a cohort of cognitively-normal older adults. Whereas a single mode was previously reported to link cognitive and behavioural traits to functional connectivity patterns within healthy adults (Smith, et al., 2015), we identified three modes capturing significant interdependencies between phenotypic measures, age and functional connectivity in our older cohort. The first mode comprises an opposition between cognitive performance and age on connectivity patterns. The second mode accounts for an independent and positive association of education with functional connectivity, whilst the third mode captures weak relations. Including age in a multivariate model of brain-behaviour relations in a healthy elder cohort thus appears to split the single mode expressed in younger adults into three separate modes, with age and education loading orthogonally.

All cognitive domains in the first mode load along a positive axis, mirroring positive traits within healthy adults (Smith, et al., 2015). Age, on the other hand, is positioned on the negative pole. This mode thus captures the opposing pull between cognitive performance and age in their co-variation with connectivity patterns. The influence of age-related changes most strongly opposes the connectivity patterns associated with greater attention and processing speed scores. The spread of other cognitive domains captured by this mode converges with the previous ageing literature: Across the lifespan, tasks assessing attention and processing speed are the most sensitive to age-related reductions in performance. Furthermore, age-related changes in lower-level abilities (i.e. perceptual speed, psychomotor abilities) are proposed to account for the reduced performance in other abilities such as memory and executive functioning (Baltes and Lindenberger, 1997; Lee, et al., 2012; Park and Reuter-Lorenz, 2009; Salthouse, 1996). The sensitivity of such sensorimotor processes is consistent with the observable slowing of daily activities in older individuals, as exemplified by mobility and driving abilities (Ball, et al., 2007). Indeed, the functional connections most positively expressed by this mode are lower-order systems linking visual and somatosensory cortices, with additional involvement of parietal association areas. These regions are connected by bilateral insular (posterior) and operculum (parietal) areas - whose functions are not only associated with simple sensorimotor tasks, but also the functional integration of sensorimotor areas (Sepulcre, 2014; Sepulcre, et al., 2012). Age-related changes observed here also build upon previously reported reductions in resting-state connectivity with age within sensorimotor systems (Betzel, et al., 2014; Chan, et al., 2014; Geerligs, et al., 2015), as well as for connectivity of the parietal operculum itself (Cao, et al., 2014; Tomasi and Volkow, 2012).

There conversely exists a network of functional connections negatively expressed by this mode involving links between pre-motor, pre- and post-central gyri and superior medial frontal areas - regions involved in planning and performing motor output (Hu, et al., 2015; Nachev, et al., 2008; Tremblay and Gracco, 2010). Whereas motor performance generally decreases with age (Ketcham and Stelmach, 2001; Seidler, et al., 2010), paradoxically increased functional activations in these areas occur during motor tasks in older subjects (Carp, et al., 2011; Heuninckx, et al., 2008; Kleerekooper, et al., 2016; Seidler, et al., 2010). Increased activation may act to compensate for changes in neural integrity (Cabeza, et al., 2002), and the decreased functional specialisation of brain regions (Seidler, et al., 2010), as reflected by the increases in between-network connectivity with age (Betzel, et al., 2014; Chan, et al., 2014; Ferreira, et al., 2015; Geerligs, et al., 2015; Grady, et al., 2016; Ng, et al., 2016). Interestingly, a longitudinal study revealed from ages between 65-70 years, an increase over time (two-year interval) for functional connectivity between the default-mode and executive networks (Ng, et al., 2016). Increases in between-network connectivity over time in their study were further associated with reductions in processing speed performance: The functional connections most negatively expressed by the first mode in the present study also largely involve the between-network interactions of default-mode and control-related areas. Hence, the first mode may capture the dynamic changes to brain connections with age (Moran, et al., 2014), whereby patterns of more efficient connectivity (relatively lower connectivity) are also associated with better (and younger) performers.

Of interest, education loads only weakly on to the networks expressed by the age-related changes in cognitive performance (i.e. mode one). The circuits supporting sensorimotor functions in older adults may thus be resistant to moderating factors such as years of education. This is in apparent contradiction to the mitigation of age-related cognitive-changes and relative maintenance of volumetric brain structure observed with CR proxies in healthy older individuals (Bartrés-Faz and Arenaza-Urquijo, 2011; Stern, 2002; Stern, 2016). Years of education is a frequently employed proxy of CR, and correlates highly with independently-derived measures of brain maintenance and CR (Habeck, et al., 2016; Steffener, et al., 2016). In our data, education instead loads upon a second mode, whose functional connections are distinct from the first mode. Connections occur between visual, salience, superior parietal, and somatomotor regions, and converge upon the lateral prefrontal areas - circuitry (especially fronto-parietal links) consistently implicated in cognitive control and other higher-order functions (Cocchi, et al., 2013; Hearne, et al., 2015; Koechlin, et al., 2003; Spreng, et al., 2010). Indeed, executive function partially loads onto the connectivity patterns expressed by this mode, revealing increased education may at least provide partial neuroprotection for tasks comprising this domain. We note that executive functions represent heterogeneous cognitive processes, as reflected by the additional components that are tapped into by the domain composites (i.e. TMT B and FAS tasks). The unique variance captured by the connectivity patterns of the first mode presumably reflects the diverse aspects of the functions they assess (and hence lower internal-consistency) (Greenaway, et al., 2009; Sanchez-Cubillo, et al., 2009). However, the two tasks do project almost identically onto the second mode: As noted, the expressed connections of this mode are consistent with those supporting executive/control-related processes.

A third mode links relatively weak positive associations between connectivity patterns to memory and visuo-spatial abilities. This third mode may capture cognitive correlates relatively independent of subject’s age and education. However, this mode was only weakly significant in our primary analysis and did not generalize to auxiliary analyses.

We observed that NART IQ scores loads with other cognitive domains in opposition to age on the first mode, while education remains independently captured by the second. This divergent loading of intelligence and education on the first mode is interesting given both measures represent typical proxies of CR (Xu, et al., 2015), are highly-correlated, and share similar co-variation with functional connectivity patterns observed in younger adults (Smith, et al., 2015). However, CR proxies have previously been shown to mitigate age-related changes independent of each other (Richards and Sacker, 2003b; Stern, et al., 1995; Suo, et al., 2012). The functional connections expressed within the second mode of this CCA are predominately between default-mode (inferior and medial parietal regions) and control-network hub-areas (middle frontal gyrus) to other task-affiliated networks. Previous research has established that higher-order cognitive functions are dependent upon by these transmodal hub-areas (Cole and Schneider, 2007; Raichle, 2015; Seghier, 2013; Utevsky, et al., 2014). The predominance of between-network interactions loading with increased education is salient given that the integration of functional subsystems is critical upon cognitively-demanding tasks (Bassett, et al., 2011; Braun, et al., 2015; Cocchi, et al., 2013). In our data, intelligence loads moderately upon the age-related networks of the first mode, whilst the influence of education is relatively strongest for non-specific between-network interactions. Despite these CR proxies being highly-interwoven, this divergence could be attributed to intelligence representing innate contributions towards normal ageing (Deary, et al., 2010; Plomin and Deary, 2015), whilst educational attainment is perhaps more reflective of modifying factors. Further investigations exploring the rich spatiotemporal structure of resting-state (Madhyastha and Grabowski, 2014; Zalesky, et al., 2014) and task-based fMRI patterns within this older cohort may disentangle the benefits of increased education upon these non-specific between-network interactions.

We did not include sex in our primary analyses, as we sought to elucidate general age-related changes across our cohort. Including participants’ sex within the CCA model allows a nested investigation of the influence of sex on age-related brain-behaviour correlates. This analysis revealed a similar latent structure of phenotypic inter-relations to the first and second mode of the original analysis. Within the first mode, sex (males) loaded onto negative associations with functional connectivity patterns, hence with age and in opposition to better cognitive performance. Here, males demonstrate poorer performance on memory-based tasks, which is consistent with the cognitive styles of males from both young and older adult populations (Gur, et al., 2012; Hoogendam, et al., 2014; Kimura, 2004). Sexual dimorphisms in brain connectivity and structure are also consistently observed across both young and older adults (Feis, et al., 2013; Ingalhalikar, et al., 2014; Joel, et al., 2015; Perry, et al., 2015). The inclusion of sex within the CCA has only minimal impact spatially on the functional edges most strongly expressed by both positive and negative associations within the first mode. Our data thus suggest that sexual dimorphisms in later life load on top of background age-related changes, particularly for the circuits supporting memory functions. In contrast, the connections most strongly expressed in the second mode are substantially influenced when including sex. We note that performance in visuospatial ability, executive functioning, and male sex share similar co-variations here with functional connectivity patterns. In the current sample males demonstrate greater education years, and hence these uniquely expressed connectivity patterns may reflect the benefits of their educational attainment for such cognitive processes (Gur, et al., 2012; Hoogendam, et al., 2014; Kimura, 2004).

The relatively large cohort and the multivariate nature of CCA bring new insights into the relationship between age, cognition and functional brain networks. However, these findings should be interpreted in light of a number of limitations. The cross-sectional and association-based nature of the study design precludes causal inferences. A formal analysis of the influence of age and the relative preservation of age-related changes with greater educational attainment would mandate a longitudinal within-subjects design (Stern, 2016). CR itself represents an inherently complex construct (Stern, 2016), with an individual’s innate ability and neuroplastic experiences contributing to the slowing of age-related changes in similar (Habeck, et al., 2016; Steffener, et al., 2016) and independent forms (Richards and Sacker, 2003a; Suo, et al., 2012). Nonetheless, education years remains one of the most widely implemented CR proxies (Bartrés-Faz and Arenaza-Urquijo, 2011; Xu, et al., 2015), and is also inextricably intertwined with the enriching lifestyle choices that individuals pursue (Ross and Wu, 1996; Valenzuela and Sachdev, 2007; Xu, et al., 2015).

The individual tests were partitioned into cognitive groupings as part of the broader longitudinal study (MAS): This was done to facilitate the longitudinal assessment of the current study participants, and to compare our findings with widely-adopted theoretical constructs. While there is no complete consensus regarding which cognitive domain particular tests should be allocated to, our choice was guided by a review of the extant literature and accorded with the widespread practice used among neuropsychologists (Lezak, Howieson, & Loring, 2004; Strauss, Sherman, & Spreen, 2006; Weintraub et al., 2009). We acknowledge neuropsychological tests are multifactorial in structure and even though a test may primarily focus on one aspect of cognition, domain performance here is indeed highly-correlated, and is thus potentially influenced by shared cognitive processes. As noted, this is particularly the case for the executive-composite. The verbal-abilities also assessed by the FAS (Controlled Oral Word Association Test) are closely related to those tapped into by the semantic fluency task, grouped within the language domain. The extant literature, however, from both healthy and clinical populations supports grouping the FAS and semantic fluency tasks into separate domains (Henry and Crawford, 2004; Schmidt, et al., 2017). FAS performance requires the suppression of semantically or associatively related words (Katzev, et al., 2013; Luo, et al., 2010; Shao, et al., 2014), and is hence typically thought to involve executive processes such as strategy, initiation, and self-monitoring (Henry and Crawford, 2004). Categorical fluency tasks (i.e. the FAS) require more cognitively-demanding resources than semantic fluency tasks (Schmidt, et al., 2017), as individuals within the latter can rely on pre-existing (sub-)categorical links to retrieve responses (Schmidt, et al., 2017). Distinct functional profiles are implicated during categorical and semantic fluency tasks (Birn, et al., 2010; Katzev, et al., 2013), along with differences in their expression with clinical populations and focal brain lesions (Henry and Crawford, 2004; Shao, et al., 2014). Semantic and categorical fluency tasks do share overlapping cognitive and neurobiological profiles, and hence cannot be considered pure assessments of a cognitive process (Henry and Crawford, 2004; Shao, et al., 2014). Nonetheless, both the TMT B and FAS primarily serve similar executive functions (Lezak, et al., 2004; Strauss, et al., 2006), and are thus grouped together here.

We additionally report the internal consistency of the domain scale-items, which provide support for our *a priori* groupings. The executive composite reports a relatively-lower scale-item homogeneity - which again is not surprising - given the multifactorial structure of such processes. We note that the use of consistency measures for two-item scales is highly contested, as they underestimate the reliability of scale-items (Eisinga, et al., 2013; Tavakol and Dennick, 2011). The alternative approach to allocating tests to domains would be to use factor analysis to form empirically-based domains based on study data. We chose not to take this approach since factors formed this way are more idiosyncratic to the cohort studied and the range of tests put into the factor analysis.

The presence of three modes in our analyses contrasts to the single mode reported in the seminal paper by Smith et al (2015). Given the older age of our cohort and the inclusion of age as a factor in our CCA, we propose that this difference reflects the influence of age on brain-behaviour correlations in later life, such that age acts independent to the potential mitigating effect of earlier education. This interpretation needs to be mindful of other differences between the studies, such as the very broad range of cognitive, lifestyle and behavioural factors in Smith et al. First, the present study inferred significance by parametric methods (i.e. Rao’s *F*), whilst Smith et al. implemented non-parametric permutations. The issue of parametric versus non-parametric model testing remains an active area of debate, with the former considered more sensitive when valid and the latter more adaptive to data set size and the nature of the distribution of the variability (Bzdok and Yeo, 2017). The current sample size is considerably smaller than the cohort of HCP participants employed by Smith et al. In our opinion, parametric inference was appropriate to ensure stable and robust linear-model fits (Bzdok and Yeo, 2017; Ghahramani, 2015). In some cases, parametric models may be more sensitive and stable than their non-parametric counterparts (Eklund, et al., 2016; Friston, 2012). It is worth noting that non-parametric models have an increasing role in multivariate fMRI analyses (Nichols and Holmes, 2002). Such models are data-driven, and unlike parametric inferences, can flexibly adapt to large data sets (Bzdok and Yeo, 2017; Ghahramani, 2015; Miller, et al., 2016).

Other study differences in Smith et al. include the use of high-temporal resolution rsfMRI data and a high dimensional ICA-based approach for de-noising and cortical parcellation. The availability of a more modestly size cohort in our study as well as differences in the characterization of our cohort precluded the application of an identical pipeline. However, the connectivity patterns expressed by the positive individual traits and behaviours of the younger population in Smith et al. are those primarily those of default-mode, control-network, and sensory-related cortices. The large-scale interactions between these areas are also particularly expressed by the higher intelligence and education levels (i.e. the second mode) of the older adults in the present study, which hence speaks to the continued contribution of positive phenotypic traits to healthy brain functioning.

In conclusion, the present study expands upon a recent multivariate analysis of behaviour and functional brain networks in young adults through extension into cognitively-normal elders. When modelling age in our elderly cohort, we find that brain-cognition relations spilt into more than one mode, with age and education loading onto separate modes of functional connectivity patterns. Age-related changes in later life are most strongly exerted upon sensorimotor networks subserving core cognitive processes such as attention and processing speed. We find that changes within these lower-level circuits are independent to moderating factors such as higher education attainment, which confer their influence independent of age-related effects. The influence of age and education here can provide an important benchmark for the study of neurodegenerative disease and furthermore has implications for behavioural interventions in elderly populations. Whereas effects of education and sex are often controlled for within ageing investigations, the present multivariate approach further highlights the rich and complex phenotypic inter-influence on functional connectivity patterns.

## Acknowledgements

This work was supported by National Health and Medical Research Council Program (350833, 118153) and the Australian Research Council (CE140100007). Address correspondence to Dr Alistair Perry, QIMR Berghofer Medical Research Institute, 300 Herston Rd, Herston QLD 4005, Australia.

## Supplementary Information | Tables and Figures

**SI Fig 1.**
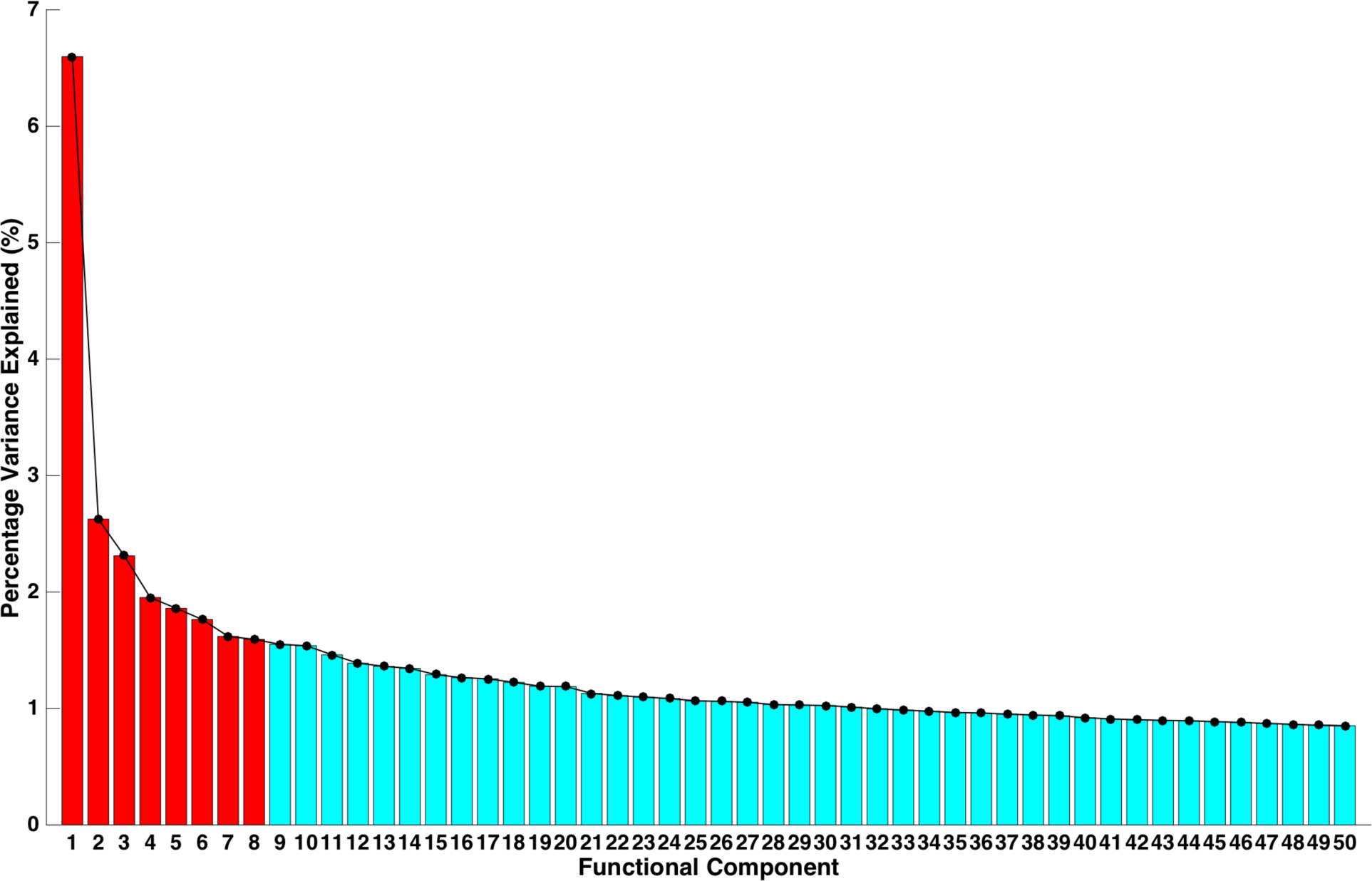
Percentage of variance explained by each principal component of the functional connectivity edges (*N_3_*). The percentage of variance explained by the first **fifty eigenvector decompositions** is shown on the *y*-axis: **The first eight eigenvectors explain 30.1% of the variance of the first 50 eigenvectors and 20.3% of the total variance.**

**SI Fig 2.**
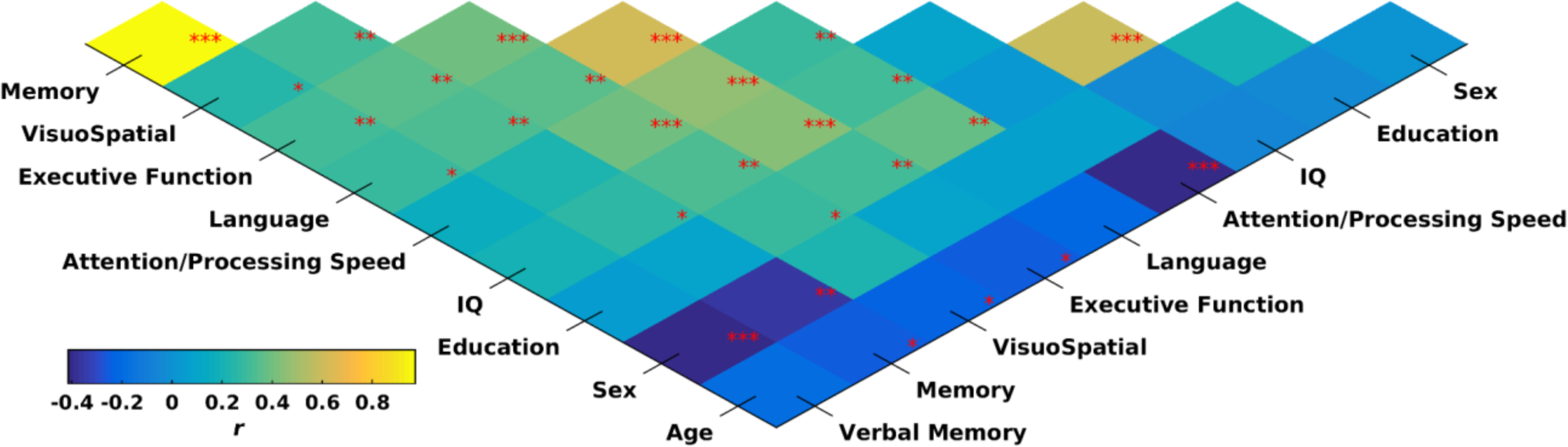
Strength and direction of relations between demographic and cognitive measures for those receiving (*n* = 91) NART IQ assessment at study baseline. * *p* < 0.05, ** *p* < 0.01, *** *p* < 0.001; FDR-corrected

**SI Fig 3.**
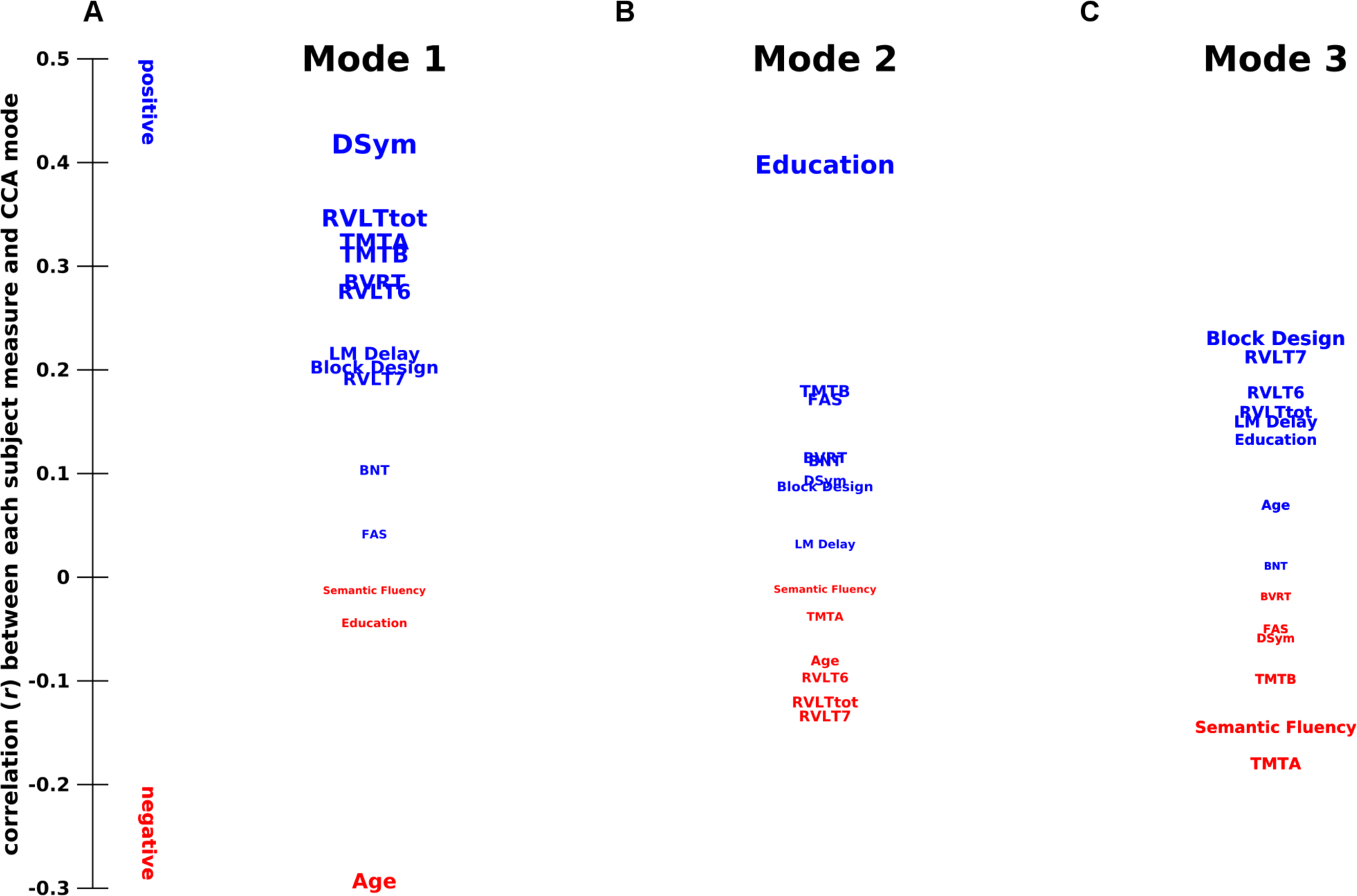
Projection of the individual neuropsychological tests onto the population co-variance captured by the original CCA. (A-C) Correlation between the individual test scores that comprise the cognitive groupings, and the functional connectivity variation (*V_m_*) captured by the original CCA. The strength and direction of the relations are indicated by vertical position and font size. DSym, Digit-Symbol Coding; TMT, Trail Making Task; RVLT, Rey Auditory Verbal Learning Test; LM delayed, Logical Memory Story A delayed recall; BVRT, Benton Visual Retention Test recognition; BNT, Boston Naming Test; FAS, Controlled Oral Word Association Test

**SI Fig 4.**
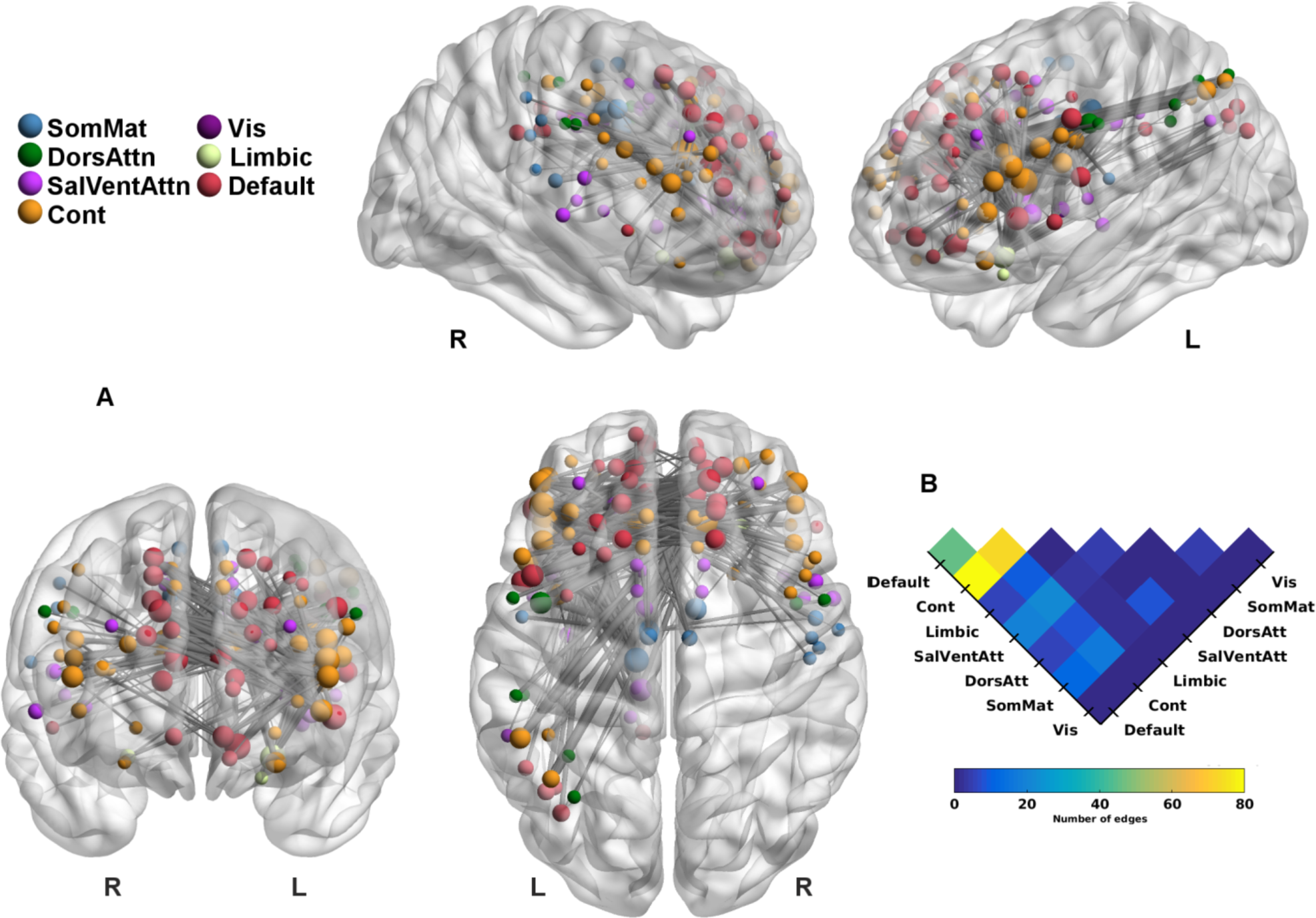
Connectivity edges most positively expressed by the third CCA mode. (A) Connectivity edges exhibiting strongest positive associations with functional connectivity patterns (*V3*). Line width indexes strength of correlation. Circle size is scaled to the number of connections each region shares within the network, whilst coloured to their functional network affiliation. The brain meshes are presented from axial (middle panel), posterior (top right), and angular perspectives of the left and right-hemisphere. (B) Coarse perspective of connectivity distributions across the functional network affiliations, with warmer colours indicating greater number of connections.

**SI Fig 5.**
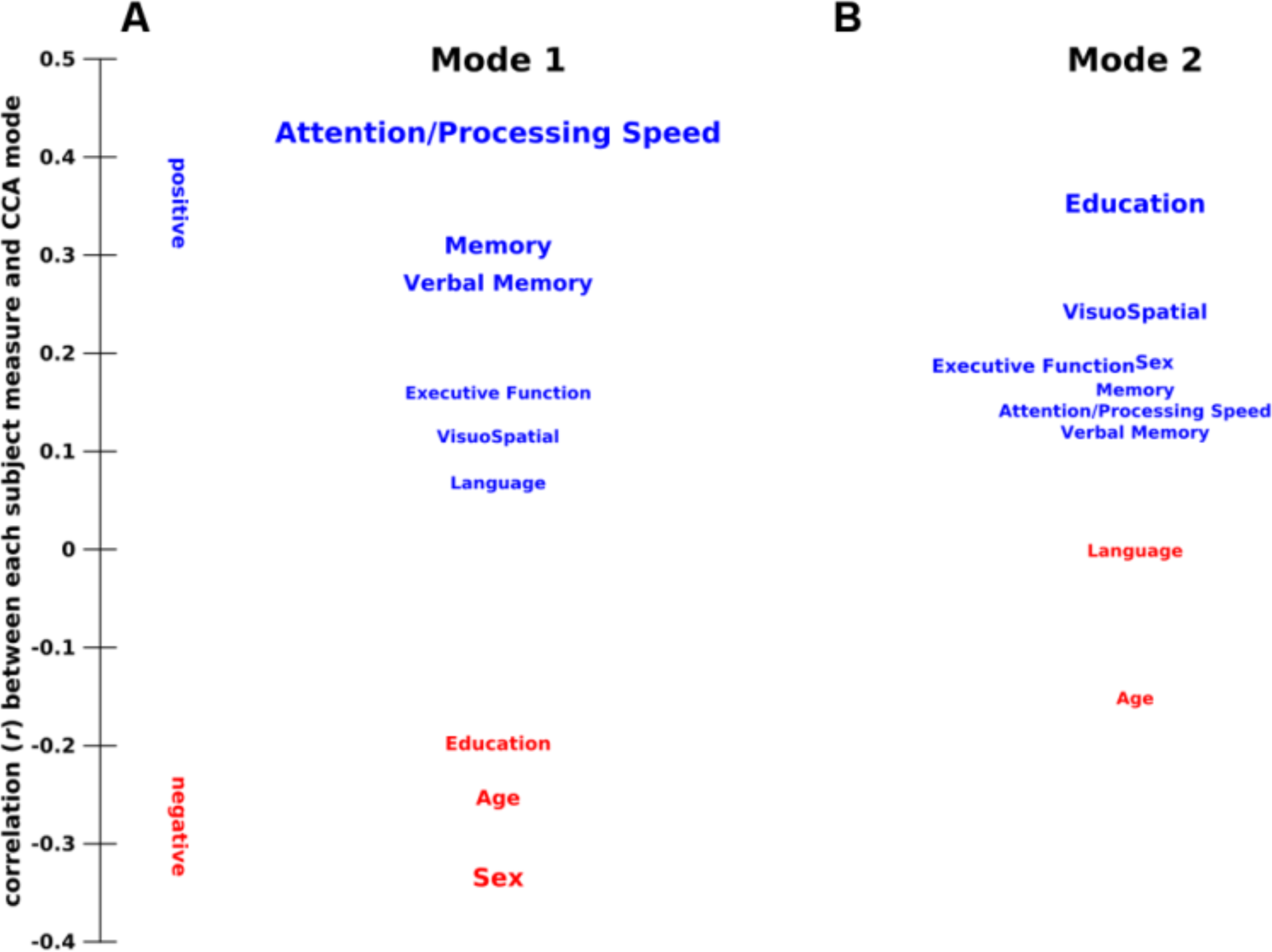
Associations between cognitive and demographic measures captured by the CCA modes (*p* < 0.05) including sex (males coded as 1). (A-B) Correlation between subject measures and functional connectivity variation (*V_m_*), with the strength and direction of the relations indicated by vertical position and font size.

**SI Fig 6.**
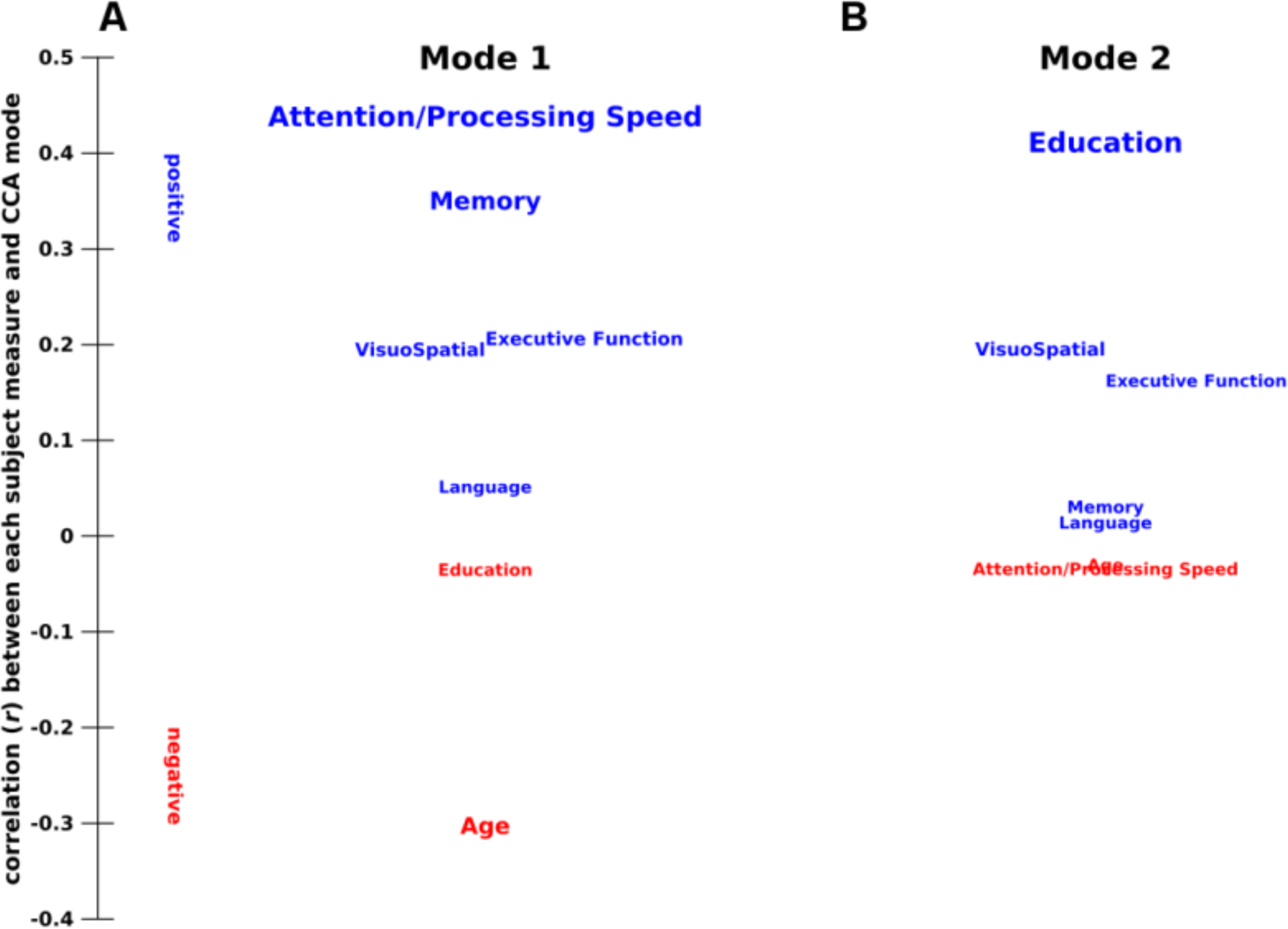
Associations between cognitive and demographic measures captured by the CCA modes (*p* < 0.05) with the removal of verbal-memory scores. (A-B) Correlation between subject measures and functional connectivity variation (*V_m_*), with the strength and direction of the relations indicated by vertical position and font size.

**SI Fig 7.**
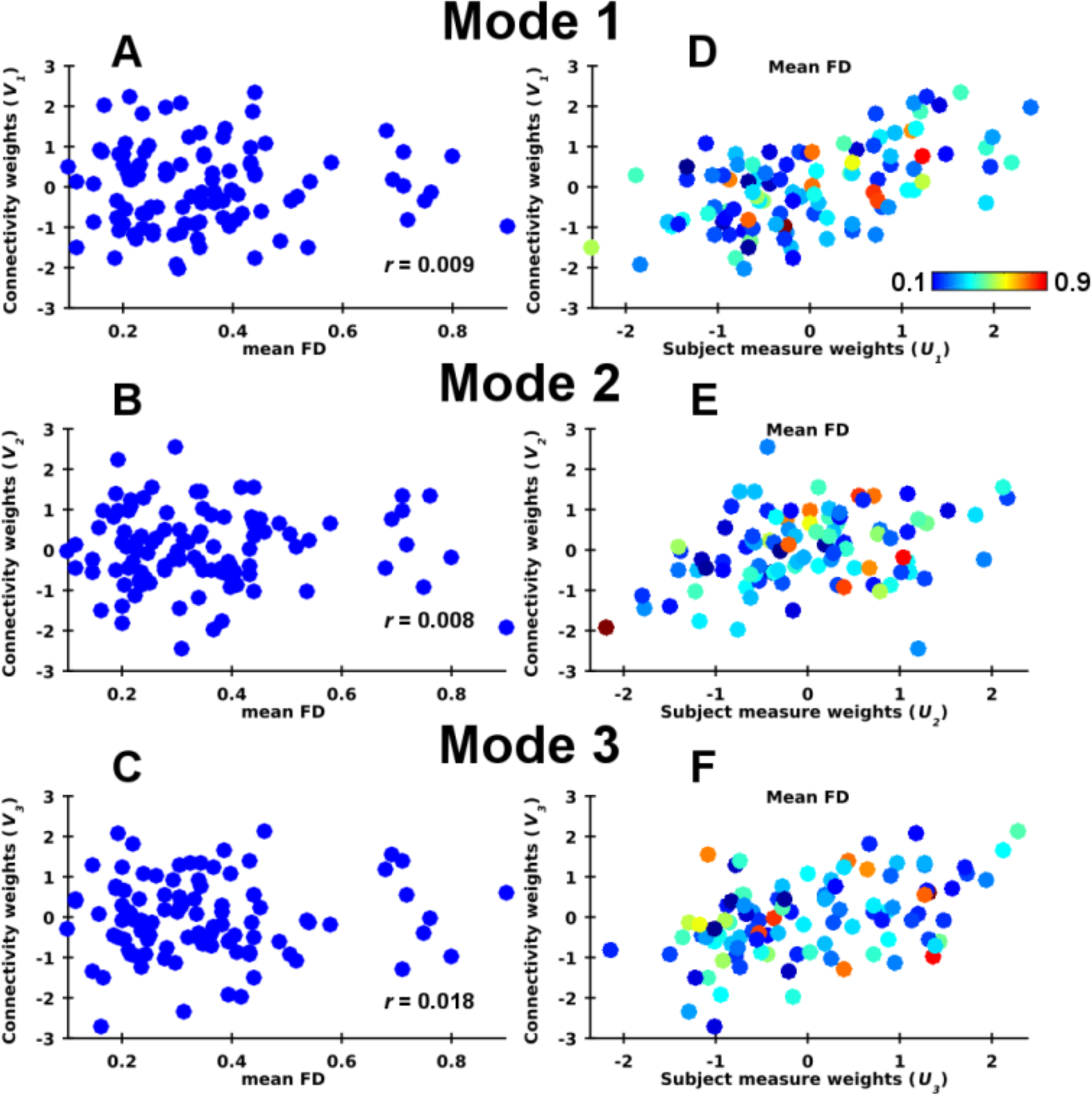
Association between subject motion and connectivity patterns across the three modes. (A-C) Scatter plots of functional connectivity patterns (*V_m_, y*-axis) as a function of mean framewise displacement (FD), showing a very weak relationship for the first (A), second (B), and third modes (C). (D-F) Each subjects weighting towards non-imaging measures (*U_m_, x*-axis) and functional connectivity patterns with the colour scaled according to subjects mean FD (cool to warm colours).

**SI Fig 8.**
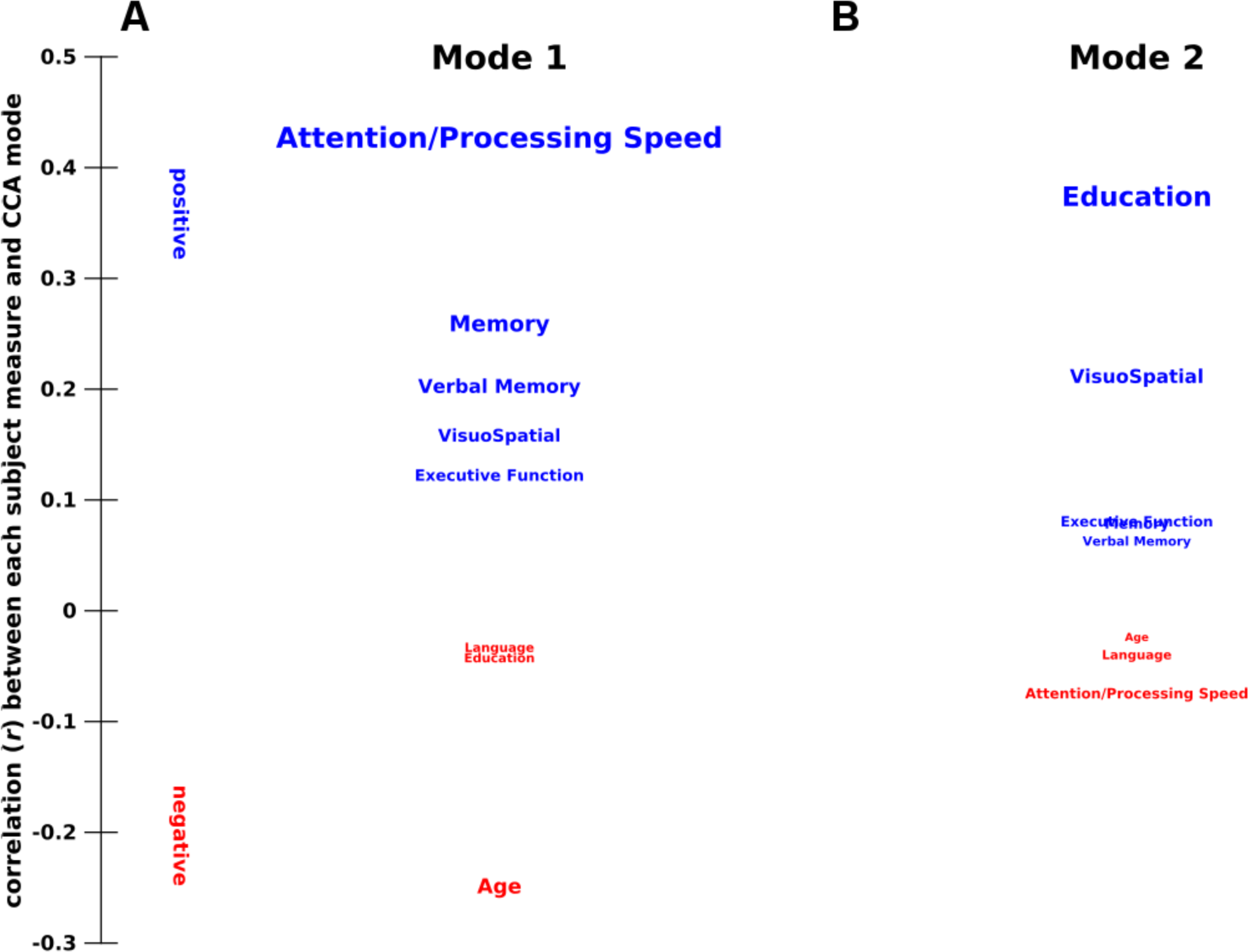
Associations between cognitive and demographic measures captured by the CCA modes(*p* < 0.05) with four functional components fed into the analysis. (A-B) Correlation between subject measures and functional connectivity variation (*V_m_*) of each mode, with the strength and direction of the relations indicated by vertical position and font size.

**SI Fig 9.**
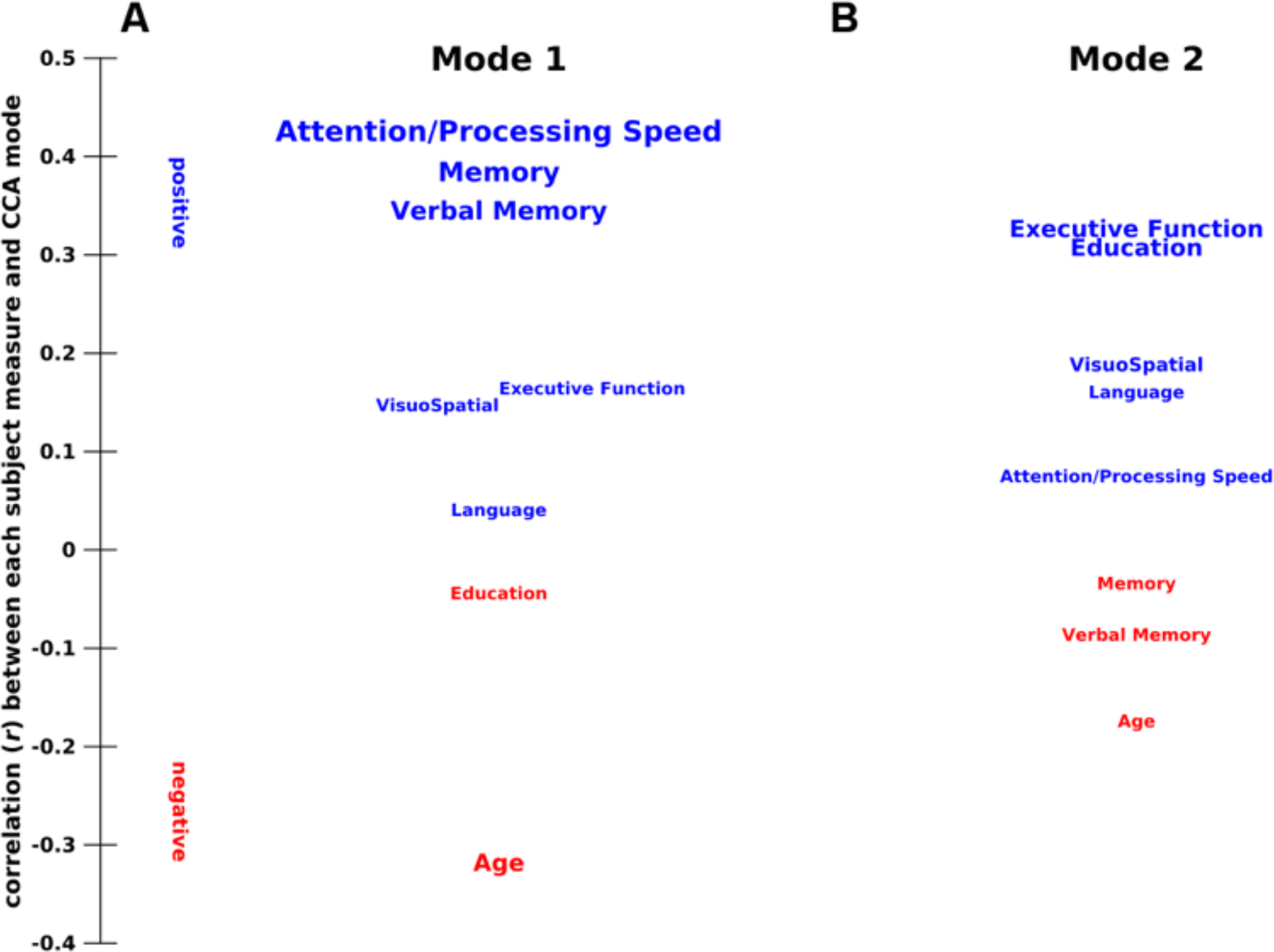
Associations between cognitive and demographic measures captured by the CCA modes (*p* < 0.05) utilizing a coarser parcellation scheme. (A-B) Correlation between subject measures and functional connectivity variation (*V_m_*), with the strength and direction of the relations indicated by vertical position and font size.

**SI Fig 10.**
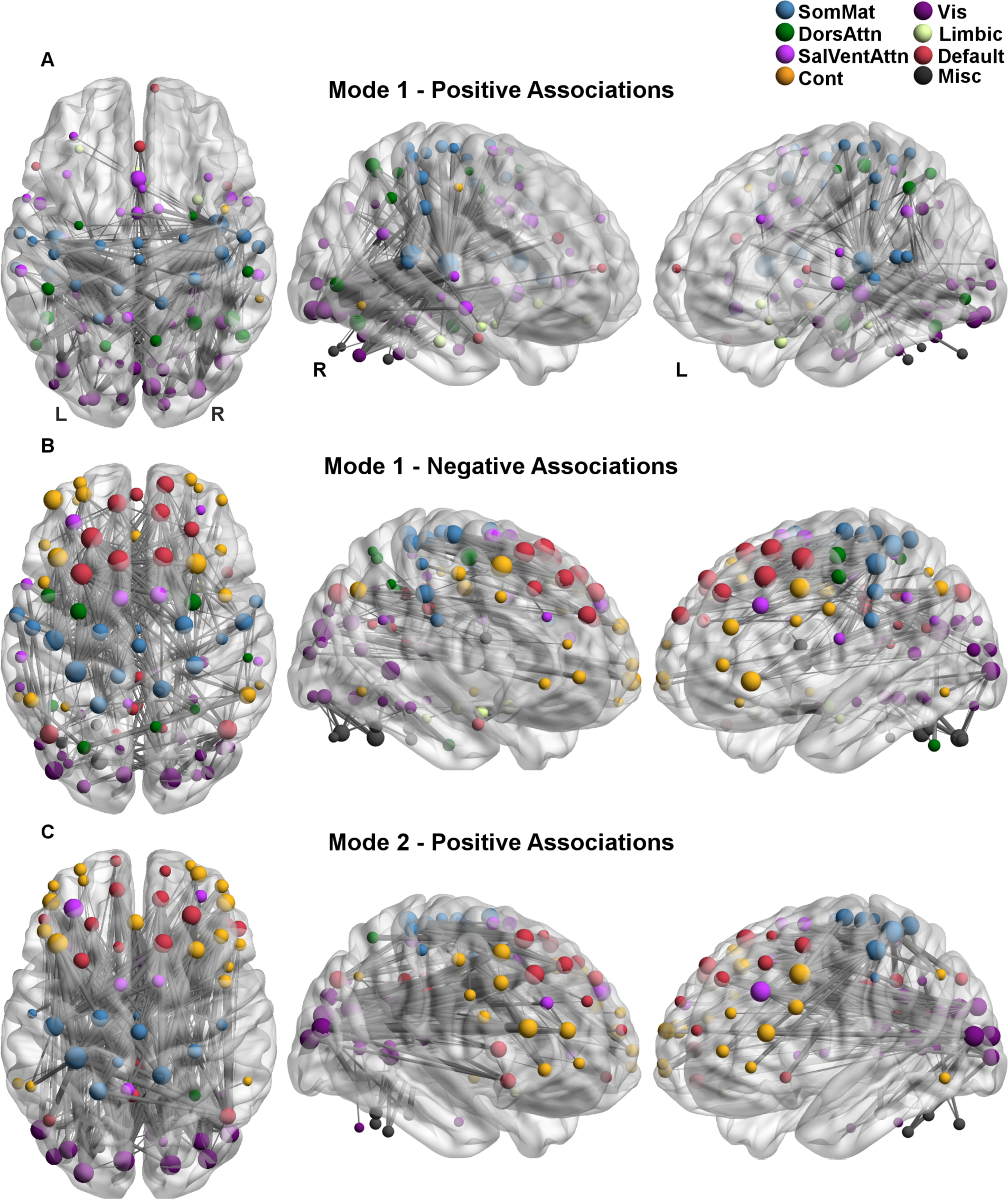
Connectivity edges most strongly expressed by the CCA modes (*p* < 0.05) with brain networks constructed by a coarser parcellation template. Connectivity edges exhibiting the strongest positive (A) and negative associations (B) with the functional connectivity patterns of the first mode (*V1*). (C) Connectivity edges exhibiting the strongest positive associations (B) with the second mode (V2). Line width indexes strength of correlation. Node size is scaled to the number of connections each region shares within the network and colour indicates their functional network affiliation. The brain meshes are presented from axial (left panel), and angular perspectives of the left (right) and right-hemisphere (middle).

**SI Fig 11.**
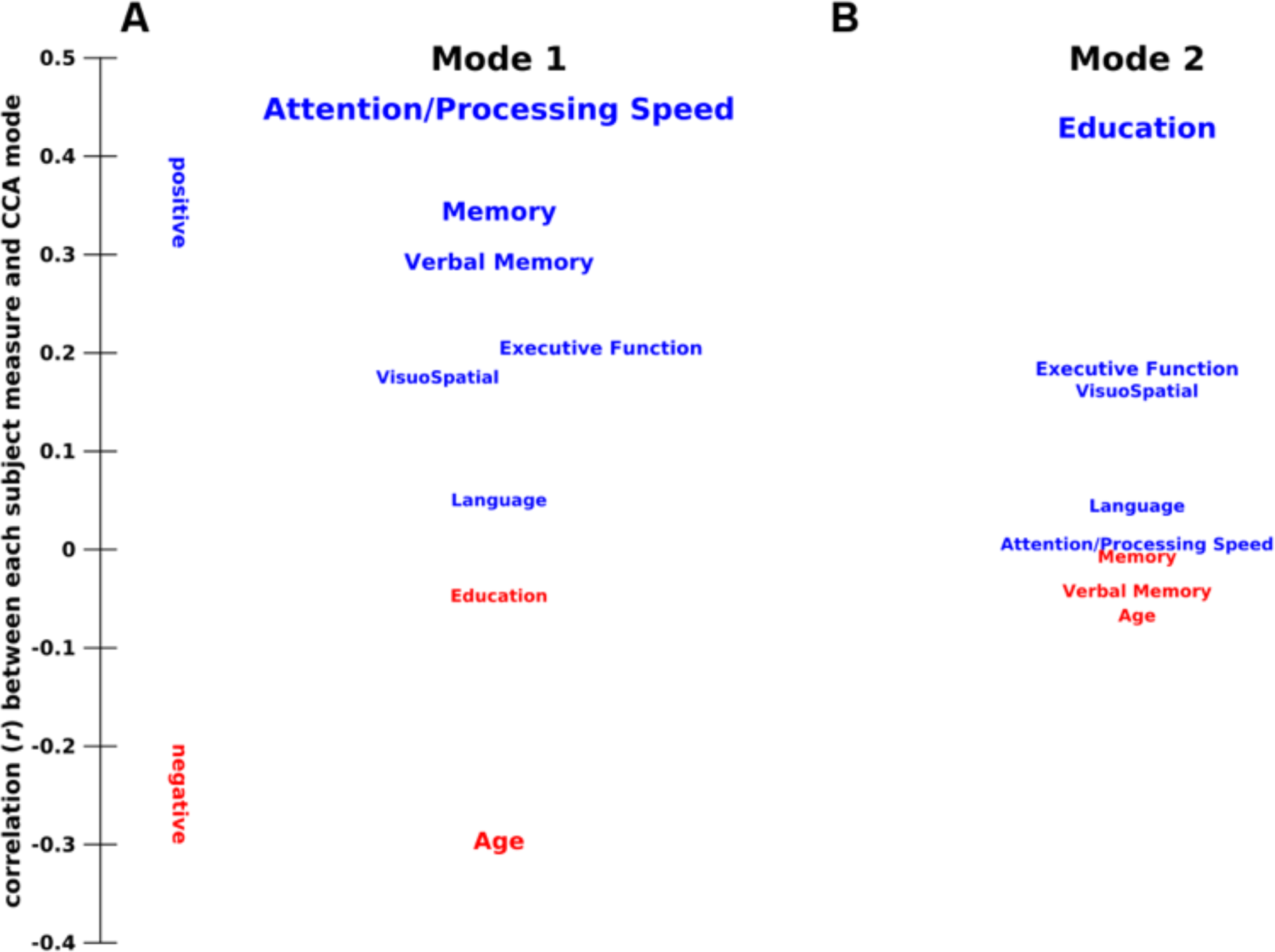
Associations between cognitive and demographic measures captured by the CCA modes (*p < 0.05*) with a smoothing kernel of 6mm applied. (A-B) Correlation between subject measures and functional connectivity variation (*V_m_*) for each mode, with the strength and direction of the relations indicated by vertical position and font size.

**SI Fig 12.**
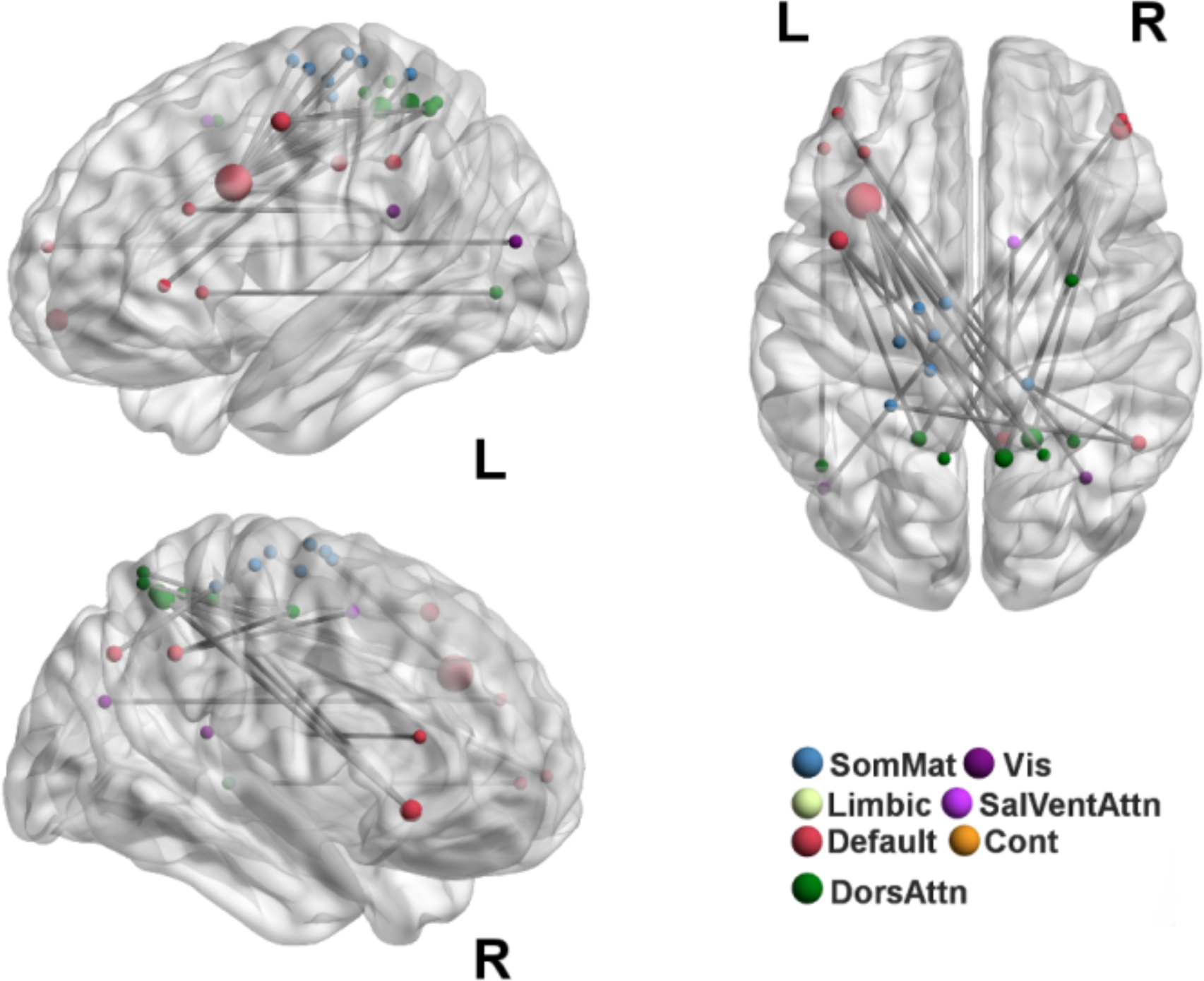
Functional connections strongly expressed for both with and without including intelligence in the second CCA mode. Circle size is scaled to the number of connections each region shares within the network, whilst coloured to their functional network specialisation. The brain meshes are presented from axial (top right panel), and angular perspectives of the left (top left) and right-hemisphere (bottom left)

**SI Table 1.** Internal consistency of cognitive domain scores.

| **Domain** | **Spearman-Brown Coefficient** |
| --- | --- |
| **Attention/Processing Speed** | 0.72 |
| **Memory** | 0.87 |
| **Verbal Memory** | 0.92 |
| **Language** | 0.65 |
| **Visuo-spatial ability** | N/A* |
| **Executive function** | 0.55 |

\* Domain composed of one neuropsychological test

**SI Table 2.** Raw performance on the individual neuropsychological tests.

| **Test** | **Mean (+- *SD*)** | **Skewness** | **Kurtosis** |
| --- | --- | --- | --- |
| DSym | 49.46 (11.60) | 0.82 | 4.08 |
| TMT A | 43.74 (13.37) | 0.63 | 3.02 |
| TMT B | 112.08 (43.52) | 1.21 | 4.52 |
| Semantic fluency | 16.84 (3.89) | 0.43 | 3.94 |
| RVLT total | 45.81 (9.28) | 0.055 | 2.55 |
| RVLT6 short-term | 9.57 (2.83) | -0.35 | 2.10 |
| RVLT7 long-term | 8.95 (3.23) | -0.25 | 2.39 |
| LM delayed | 11.36 (3.08) | 0.36 | 2.93 |
| BVRT | 12.67 (1.51) | -0.39 | 2.75 |
| BNT | 26.43 (2.53) | -0.76 | 3.40 |
| FAS | 40.46 (12.39) | 0.16 | 2.76 |
| Block Design | 24.64 (7.40) | 0.47 | 2.69 |

DSym, Digit-Symbol Coding; TMT, Trail Making Task; RVLT, Rey Auditory Verbal Learning Test; LM delayed, Logical Memory Story A delayed recall; BVRT, Benton Visual Retention Test recognition; BNT, Boston Naming Test; FAS, Controlled Oral Word Association Test

**SI Table 3.**
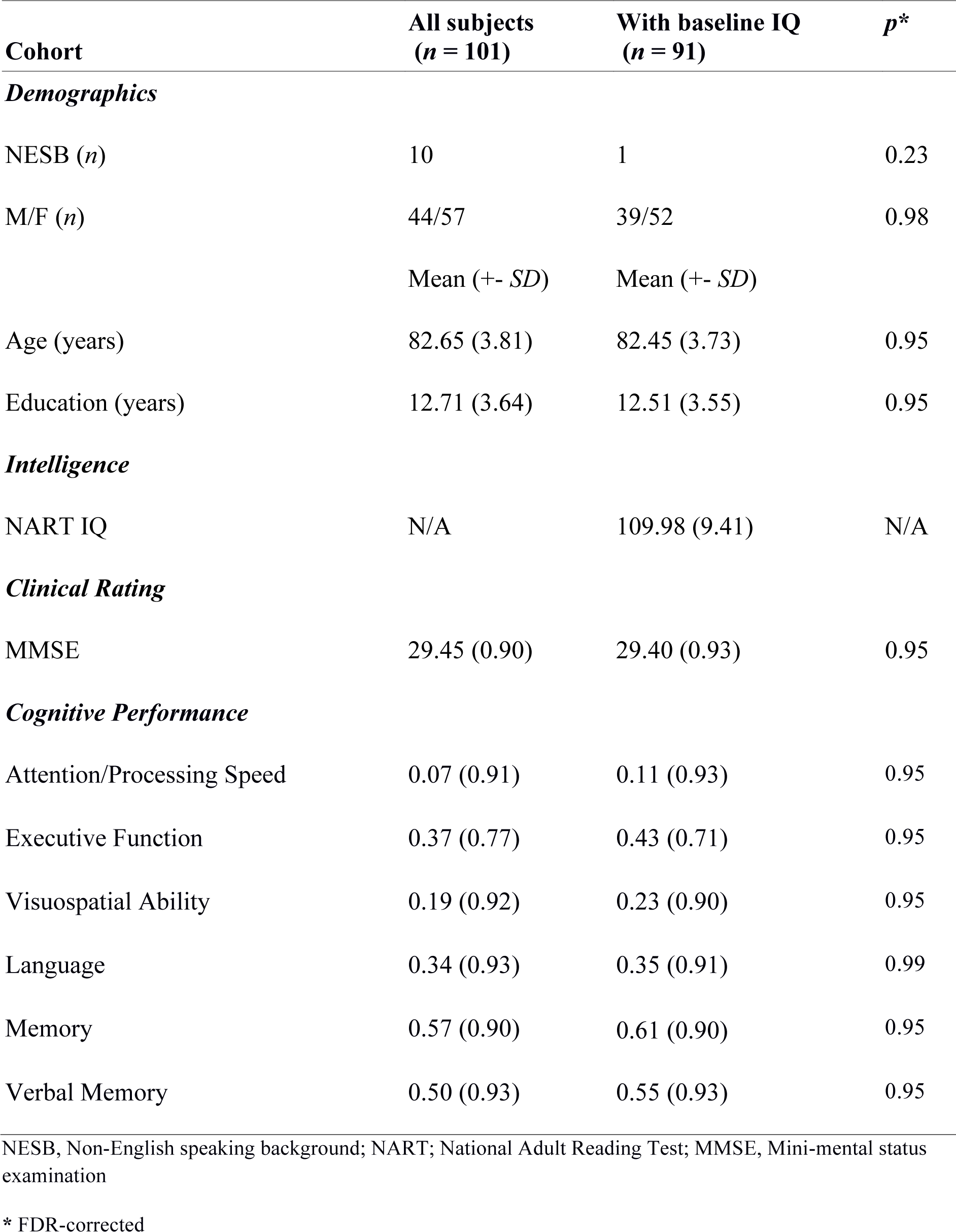
Phenotypic information for both the full sample and population subset.

**SI Table 4.** CCA modes (*p* < 0.05) with the inclusion of sex.

| CCA Mode | One | Two |
| --- | --- | --- |
| $df_1$ | 81 | 64 |
| $df_2$ | 545.36 | 490.99 |
| $F$ | 1.64 | 1.38 |
| $\lambda$ | 0.24 | 0.38 |
| $R^2$ | 0.36 | 0.25 |
| RI | 0.067 | 0.038 |
| $p$ | 0.00075 | 0.033 |

$\lambda$ = Wilk's lambda; RI = redundancy index

**SI Table 5.** CCA modes (*p* < 0.05) with including intelligence.

| **CCA Mode** | **One** | **Two** |
| --- | --- | --- |
| $df_1$ | 81 | 64 |
| $df_2$ | 480.73 | 433.31 |
| $F$ | 1.65 | 1.38 |
| $\lambda$ | 0.21 | 0.34 |
| $R^2$ | 0.40 | 0.28 |
| **RI** | 0.077 | 0.038 |
| $p$ | 0.0008 | 0.037 |

$\lambda$ = Wilk's lambda; RI = redundancy index

**SI Table 6.** CCA modes (*p* < 0.05) with the removal of verbal-memory scores.

$\lambda$ = Wilk's lambda; RI = redundancy index **SI Table 6.** CCA modes ( $p < 0.05$ ) with the removal of verbal-memory scores
| CCA Mode | One | Two |
| --- | --- | --- |
| $df_1$ | 49 | 36 |
| $df_2$ | 446.11 | 389.20 |
| $F$ | 1.99 | 1.68 |
| $\lambda$ | 0.37 | 0.53 |
| $R^2$ | 0.31 | 0.21 |
| RI | 0.044 | 0.030 |
| $p$ | 0.00017 | 0.0097 |

$\lambda$ = Wilk's lambda; RI = redundancy index

**SI Table 7.** CCA modes (*p* < 0.05) with four functional eigenvectors.

| **CCA Mode** | **One** | **Two** |
| --- | --- | --- |
| $df_1$ | 32 | 21 |
| $df_2$ | 329.81 | 258.98 |
| $F$ | 2.29 | 1.80 |
| $\lambda$ | 0.48 | 0.68 |
| $R^2$ | 0.30 | 0.21 |
| **RI** | 0.049 | 0.026 |
| $p$ | 0.00015 | 0.019 |

$\lambda$ = Wilk's lambda; RI = redundancy index

**SI Table 9.**
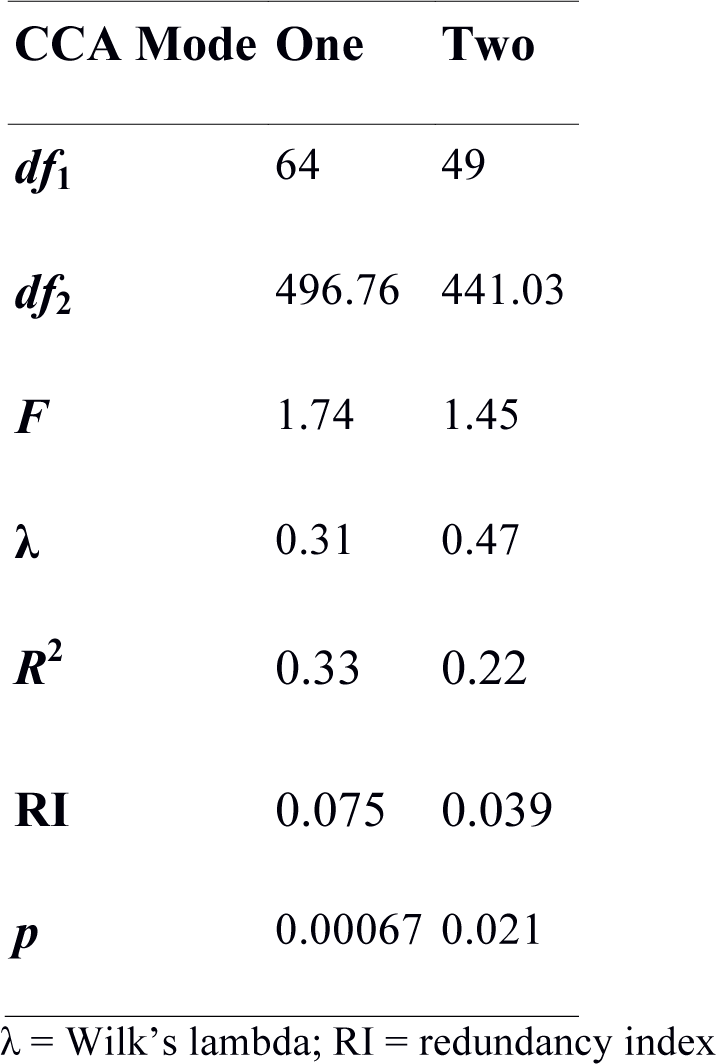
CCA modes (*p* < 0.05) with a coarser parcellation scheme.

| **CCA Mode** | **One** | **Two** |
| --- | --- | --- |
| $df_1$ | 64 | 49 |
| $df_2$ | 496.76 | 441.03 |
| $F$ | 1.74 | 1.45 |
| $\lambda$ | 0.31 | 0.47 |
| $R^2$ | 0.33 | 0.22 |
| **RI** | 0.075 | 0.039 |
| $p$ | 0.00067 | 0.021 |

$\lambda$ = Wilk's lambda; RI = redundancy index

**SI Table 10.** CCA modes (*p* < 0.05) with a smoothing kernel of 6m.

| CCA Mode | One | Two |
| --- | --- | --- |
| $df_1$ | 64 | 49 |
| $df_2$ | 496.76 | 441.03 |
| $F$ | 1.75 | 1.52 |
| $\lambda$ | 0.30 | 0.45 |
| $R^2$ | 0.32 | 0.22 |
| RI | 0.071 | 0.032 |
| $p$ | 0.00055 | 0.017 |

$\lambda$ = Wilk's lambda; RI = redundancy index

